# Mechanism of ERK-mediated Rho Activation and Stress Fiber Assembly for Cell Migration

**DOI:** 10.1101/2025.11.15.688645

**Authors:** Akib M. Khan, Jakaria Shawon, Jared P. Bergman, Keith R. Carney, Michelle C. Mendoza

## Abstract

The growth factor-activated RAS/Extracellular Regulated Kinase (ERK) pathway is a fundamental regulatory pathway that induces cell migration. ERK supports protrusion of a leading edge and also activates the small GTPase Rho, which induces actin assembly into contractile stress fibers that pull the cell body forward. Yet, the mechanism behind ERK’s induction of Rho in the cell body has remained elusive. We discover here that ERK controls Rho activity and contractile stress fibers by inhibiting Ezrin, a protein that physically links proximal actin filaments to the plasma membrane, but which also inhibits Rho by recruiting and activating ARHGAP18. ERK specifically reduces Ezrin activity in the cell body by phosphorylating the C-terminal tail of the Ezrin-activating kinase lymphocyte-oriented kinase (LOK). This phosphorylation inhibits LOK’s activation of Ezrin, thereby releases Ezrin’s inhibition of Rho and stress fibers. The ERK-LOK-Ezrin-ARHGAP18-Rho signal provides key mechanistic insight into how ERK, activated during development, wound healing, and cancer, induces Rho activity and stress fibers for cell migration.

## Introduction

Mesenchymal-mode cell migration is essential to development, immune response, and cancer cell dissemination. This manner of cell movement is realized through the integrated motion of leading-edge protrusion, cell body contraction, and rear retraction (*1*). Protrusions are driven by actin polymerization against the membrane and the assembly and disassembly of adhesions, molecular assemblies that transmit traction force between the actin cytoskeleton and cell substrate (*2*). In the cell body, contractile actin filaments called stress fibers physically squeeze and pull the cell forward (*1*, *3*). The level of actin fiber assembly and contraction, dictated by the Rho family small GTPases, is a critical determinant a cell’s ability to move (*1*, *4*). Insufficient Rho and the resulting loss of contractile fibers immobilizes cells with flattened, uniformly protruded shape (*5*). In contrast, high Rho and stress fiber assembly elongates cells with polarized morphology (*6*). Even greater, excessive Rho can constrain protrusion and contract the cell cortex, causing cells to de-adhere from the substrate and round up (*6*, *7*). Thus, balanced and properly-localized Rho activity and stress fiber assembly promotes migration (*1*, *8*).

The growth factor-activated RAS→RAF→MEK→ERK pathway is one of the principle signaling pathways that controls leading edge protrusion and cell body contraction for cell migration (*9*). The effector kinase Extracellular Regulated Kinase (ERK) is responsible for migration-driven pathogenesis in developmental disorders and cancer metastasis (*10*). Current understanding of ERK’s migration control concludes that ERK acts through Rho isoforms in leading edge protrusions and in the cell body, where the GTPases promotes polarized actin assembly and actomyosin contraction, respectively (*11–14*). In cells with oncogenic or gain-of-function ERK activation, further activation of ERK with growth factors is followed by and required for Rho activation in the leading edge (*12*). In collectively migrating cell sheets, protrusion-mediated ERK activation is followed by and required for activation of the Rho effector kinase Rho-associated kinase (ROCK) and cell contraction (*13*, *14*). Sustained RAS and ERK activation can also induce epithelial-to-mesenchymal transition (EMT), with associated Rho and stress fiber activation and cell migration (*15*, *16*). Yet, despite decades of study, how ERK controls Rho activity for pro-migratory stress fiber assembly has remained elusive.

The stress fibers responsible for cell body contraction, and their regulation by Rho isoforms, are distinct from the assemblies in leading edge protrusion (*17*), suggesting independent mechanisms of control. In leading edge protrusions, transverse arcs support the generation of pushing force that moves the membrane outward and dorsal fibers connect to and participate in the maturation of focal adhesions (*18–20*). In the cell body, contractile fibers, also called ventral stress fibers, are bundled with myosin II motor proteins and anchored to mature focal adhesions (*21–23*). Myosin-mediated contraction of ventral stress fibers generates the force that pulls the back of the cell forward during migration (*21–23*). Both RhoA and RhoC family members function in leading edge protrusions and RhoA is also necessary and sufficient for the formation and contraction of stress fibers in the cell body and rear (*24*). A population of RhoA acts in protrusions to activate the formin mDia1 to build actin filaments that support new adhesions and transverse arcs (*25–28*), while RhoC acts at the protrusion base to limit cofilin activity, thereby restricting the actin pool used to generate protrusions and conferring polarized, directional movement (*26*). In the cell center, RhoA activates Rho kinase (ROCK), which activates myosin II for stress fiber contraction to pull the cell body forward (*29*, *30*). While transverse arcs and dorsal stress fibers can be converted into ventral stress fibers (*20*, *31*), the requirement for RhoA in the formation and contraction of stress fibers in the cell body suggests that ERK may induce cell body stress fibers by activating RhoA.

Rho GTPases are active when bound to GTP, such that localized activation Rho GTPase exchange factors (GEFs) and GTPase activating proteins (GAPs) create pools of active and inactive Rho, respectively (*32*). Reports on biochemical interactions between ERK, Rho, and Rho GEFs and GAPs have not identified a plausible mechanism by which ERK controls Rho-generated ventral stress fibers in the cell body. While ERK can directly phosphorylates Rho S88 and T100 *in vitro*, which increases RhoA-GTP levels, how this phosphorylation might generate localized Rho activation in polarized cells is unclear (*33*). ERK also phosphorylates GEF-H1 at T678 to activate Rho and at S959 to inhibit Rho (*34*, *35*). While neither phospho-site has been tested for a role in cell migration, GEF-H1 primarily localizes to and is responsible for generating active Rho leading edge protrusions (*36*). ERK has also been found to phosphorylate p190GAP, which inactivates both RhoA and RhoC (*37–39*). p190GAP localizes to new adhesion sites in leading edge protrusions and the cell edge, promoting protrusion and cell polarization (*26*, *40*, *41*). We hypothesized that ERK’s control of Rho and stress fibers for migration might be mediated by ERK regulation of a RhoA-specific GEF or GAP that localizes to the cell body to such as ARHGAP18 (*42*, *43*), rather than established GEFs or GAPs involved in protrusion. ARHGAP18 is diffusedly localized in cytoplasm, where it inhibits stress fiber formation and focal adhesions (*42*). ARHGAP18 localization and activation is controlled by interaction with the Ezrin member of the Ezrin, Radixin, Moesin (ERM) family, linker proteins that connect proximal actin filaments to the plasma membrane (*44*, *45*). Ezrin is typically the most highly expressed member of the ERM family in epithelial and cancer cells (*46–48*). When Ezrin is in an open, active conformation, its FERM domain binds ARHGAP18 which enhances its GAP activity towards Rho (*45*, *49*).

Here, we test the role of these putative ERK-Rho signaling mechanisms in ERK’s induction of contractile stress fibers and cell migration. Using migratory cancer cell lines with activated ERK, we discover that Ezrin inhibits Rho and stress fibers in the cell body and that ERK induces Rho and cell migration by phosphorylating the Ezrin kinase LOK to inhibit Ezrin.

## Results

### ERK promotes contractile stress fibers and Rho activity through ARHGAP18

We characterized the ERK-controlled stress fibers during migration using H1299 metastatic lung cancer cells, which harbor an activating mutation in N-RAS that results in hyperactive ERK. We plated the cells at low density and stained for polymerized actin during active migration using phalloidin. In control, DMSO-treated cells, confocal imaging of the ventral cell area revealed stress fibers oriented parallel with the elongated cell axis (Figure 1A). Treatment with both ERK (SCH772984) and Rho (C3 exoenzyme) inhibitors depleted the ventral stress fibers. Transfection of mCherry-tagged myosin regulatory light chain 2 (MLC) showed that myosin II co-localized with the stress fibers, confirming their contractile nature (Figure 1B and 1C). ERK inhibitor reduced stress fiber co-localization with myosin II (Figures 1B and 1C). We quantified stress fiber length within the cell body, since the length of thick actin filaments dictates their formation of contractile units with tension (*50*). We applied hysteresis thresholding to capture the bundled actin filaments and created a cell body region of interest (ROI) for analysis by shrinking the segmented cell edge by 5 μm (Figure S1A). The actin filaments that remained under ERK and Rho inhibition exhibited a mesh-like architecture and were significantly shorter in length (Figures 1A and D), consistent with ERK functioning to build contractile stress fibers in the cell body.

**Figure 1.**
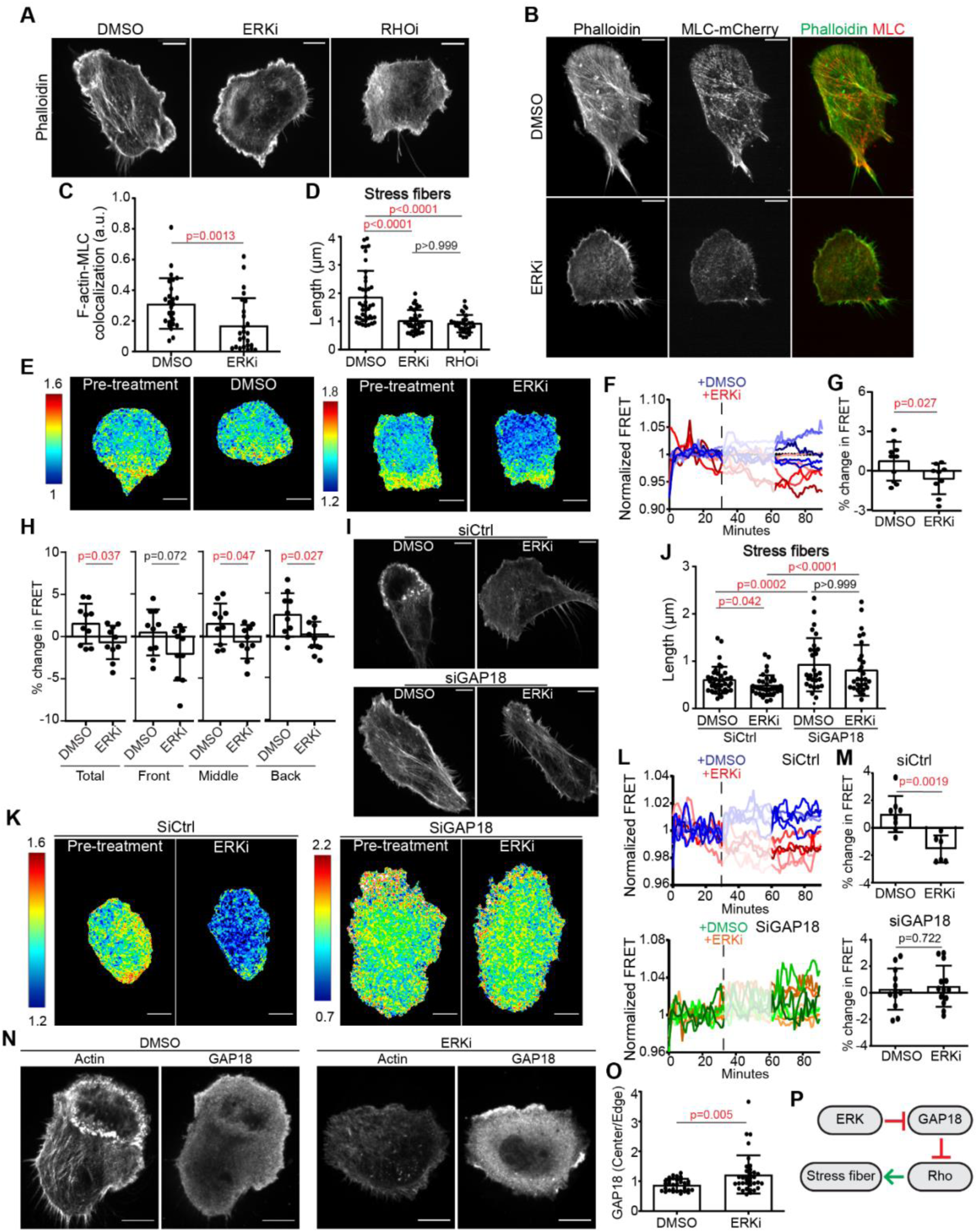
ERK promotes Rho activity and stress fiber formation through ARHGAP18. H1299 cells. (A) Phalloidin-Alexa 488 staining of cells treated with DMSO (n=43), ERK inhibitor (ERKi, SCH772984 5 µM; n=33 cells), or RHO inhibitor (RHOi, C3 exoenzyme 1 µg/mL; n=35) or with (B) expression of myosin light chain (MLC)-mCherry (DMSO, n=26; ERKi, n=25). (C) Colocalization coefficient of actin and myosin light chain. (D) Quantification of stress fiber length from A. (E) Representative Rho activity heatmaps from RhoA2G biosensor and (F) time course of activity in cells migrating under DMSO (n=10) or ERKi (n=10) treatment. FRET ratios are normalized to the mean of pretreatment values. Dashed line indicates treatment, and only the last 30 min post-treatment were used for quantification. (G, H) Quantification of percentage change in FRET ratios overall and in the defined front, middle, and back of the cell. (I, J) Representative images of phalloidin staining and quantification of stress fiber length in the cell body in cells transfected with siCtrl or siARHGAP18 and treated with DMSO or ERKi (siCtrl: DMSO n=37, ERKi n=28; siARHGAP18: DMSO n=28, ERKi n=31). (K) Representative RhoA2G activity heatmaps (L) and time courses in siCtrl and siARHGAP18 cells, before and after ERKi treatment. (M) Quantification of percentage change in FRET ratios from K, L. and Fig. S1L (siCtrl DMSO n=9, ERKi n=7; siARHGAP18: DMSO n=12, ERKi n=12). (N) Phalloidin staining and immunofluorescence for endogenous ARHGAP18 in DMSO (n=32) and ERKi-treated cells (n=36). (O) Quantification of ARHGAP18 intensity in the cell center versus edge from N. (P) Schematic of ERK signaling for Rho activation and stress fiber formation. Scale bars 10 µm. Significance by Student’s t-test (C, G, M, O) or one-way ANOVA (D, H, J).

We next tested if ERK regulates a cell body-localized pool of RhoA that generates the contractile stress fibers. As reported previously in other cell types (*33*, *35*), ERK inhibition reduced total Rho-GTP levels assayed with the Rho-GTP binding domain of Rhotekin, (Figure S1B). We then monitored the spatial distribution of RhoA activity in migrating H1299 cells using the RhoA2G FRET biosensor, which reports an increase in FRET/CFP ratio when bound to RhoA-GTP versus RhoA-GDP (*51*). As expected, RhoA activity was present at the leading edge and in the posterior portion of the cell body and rear (Figure 1E, Supplemental movie 1). We tracked cell movement over the course of the timelapse to identify the direction of motion and divided the cell into fourths with the leading edge defined as the first fourth, the cell body as the middle half, and the cell rear as the last fourth, for each time point (Figure S1C), which confirmed that most of the active Rho localized to the posterior cell body and rear (Figure S1D). Next, we sought to identify the population of RhoA that is regulated by ERK in migrating cells. Averaging the mean cellular FRET for 30 mins before and after the addition of DMSO or ERK inhibitor showed that ERK inhibition reduced the mean Rho activity (Figures 1E-G, Supplemental movie 2). Quantifying the change in FRET in the subcellular regions showed that ERK inhibition reduced Rho activity predominantly in the middle and rear of the cell (Figure 1H), consistent with ERK inducing RhoA-mediated contractile fiber activity in the migrating cell body. Because Rho activity pulses about every ∼1 hr in collectively migrating cells (*13*, *52*), we tested if ERK also controls RhoA activity pulses in randomly migrating single cells. Quantifying the peaks of RhoA activity within the cell rear over time identified pulses that lasted ∼13 mins, with some shorter (∼7 min) and longer (∼22 min) events (Figure S1E). The standard deviation of the FRET signal, which reflects pulse frequency and amplitude, remained unchanged upon ERK inhibition at the back of the cell (Figures S1F). Thus, during single-cell migration, ERK regulates baseline RhoA activity, rather than dynamic Rho pulses.

We tested the involvement of the putative ERK-controlled Rho GEFs and GAPs in contractile fiber maintenance by knocking down GEF-H1, p190RhoGAP, and ARHGAP18 in migrating H1299 cells. Western blotting confirmed that siRNA transfection reduced total protein by 70-80% (Figures S1G-I). Knockdown of GEF-H1 or p190RhoGAP did not affect the cell body-localized contractile stress fibers, but ARHGAP18 knockdown increased the stress fiber length (Figures S1J and S1K). We further tested if ARHGAP18 mediates ERK regulation of contractile fiber activity by treating the ARHGAP18 depleted cells with ERK inhibitor. While ERK inhibition reduced actin fiber length in control cells, it did not affect fiber length in ARHGAP18-depleted cells (Figures 1I and 1J). Similarly, ERK inhibition did not affect RhoA activity in ARHGAP18 knockdown cells (Figures 1K-M and S1L). This suggested that ERK controls stress fiber assembly through ARHGAP18 and RhoA. Since Rhos are controlled by GEF and GAP subcellular re-localization, we tested if ERK controls ARHGAP18 localization to the cell body. Immunofluorescence staining for ARHGAP18 showed that ERK inhibition increased the ratio of ARHGAP18 in the cell center versus cell edge (Figures 1N and 1O). This suggested that ERK may promote cell body RhoA activity and contractile stress fibers by reducing ARHGAP18 recruitment to the cell body (Figure 1P).

### ERK inhibits Ezrin activation in the cell body

Since active Ezrin binds and controls the localization of ARHGAP18 to locally reduce Rho activity(*45*, *49*), we tested if ERK modulates Ezrin. Western blotting of subconfluent H1299 cells showed that treatment with MEK (AZD6244) or ERK (SCH772984) inhibitor increased the active, phosphorylated form of Ezrin, detected as phospho-Ezrin/Radixin/Moesin (pERM) Thr567 (Figure 2A). Two-color imaging of pERM and total Ezrin confirmed that pEzrin is the predominant pERM family member in H1299 cells and is induced by ERK (Figure S2A). MEK and ERK inhibition also increased pERM levels in second metastatic cell line with oncogenic RAS activation, A375m1 cells, indicating that ERK generally inhibits Ezrin activation (Figure 2B). We tested if ERK preferentially regulates Ezrin phosphorylation in the cell body by staining cells for pERM and epifluorescence imaging and quantification of the pERM intensity in the cell body versus cell edge. Consistent with ERK’s regulation of Rho and stress fibers, ERK inhibition increased phospho-Ezrin (pEzrin) specifically in the cell body, not at the cell edge (Figure 2D). Further, ERK-controlled Ezrin activity was restricted to the cell body when the ventral slice was imaged by confocal microscopy, in cells co-treated with PLX-4032 (a non-canonical BRAF activator) and ERK inhibitor (Figures S2B and S2C). We also tested the spatial specificity of ERK’s regulation of Ezrin utilizing the membrane-targeted Ezrin BRET biosensor (MyrPB-Ezrin-LucII and rGFP-CAAX), in which inactive, closed conformation MyrPB-Ezrin-LucII undergoes resonance energy transfer with the membrane-bound GFP(*53*). The BRET in H1299 cells treated with ERK inhibitor was indistinguishable from that in DMSO-treated controls, while cells treated with the kinase inhibitor staurosporine or phosphatase inhibitor calyculin A showed the increased (inactive Ezrin) or decreased (active Ezrin) BRET, as expected (Figure S2D). Thus, ERK primarily inhibits Ezrin activity in the cell body.

**Figure 2.**
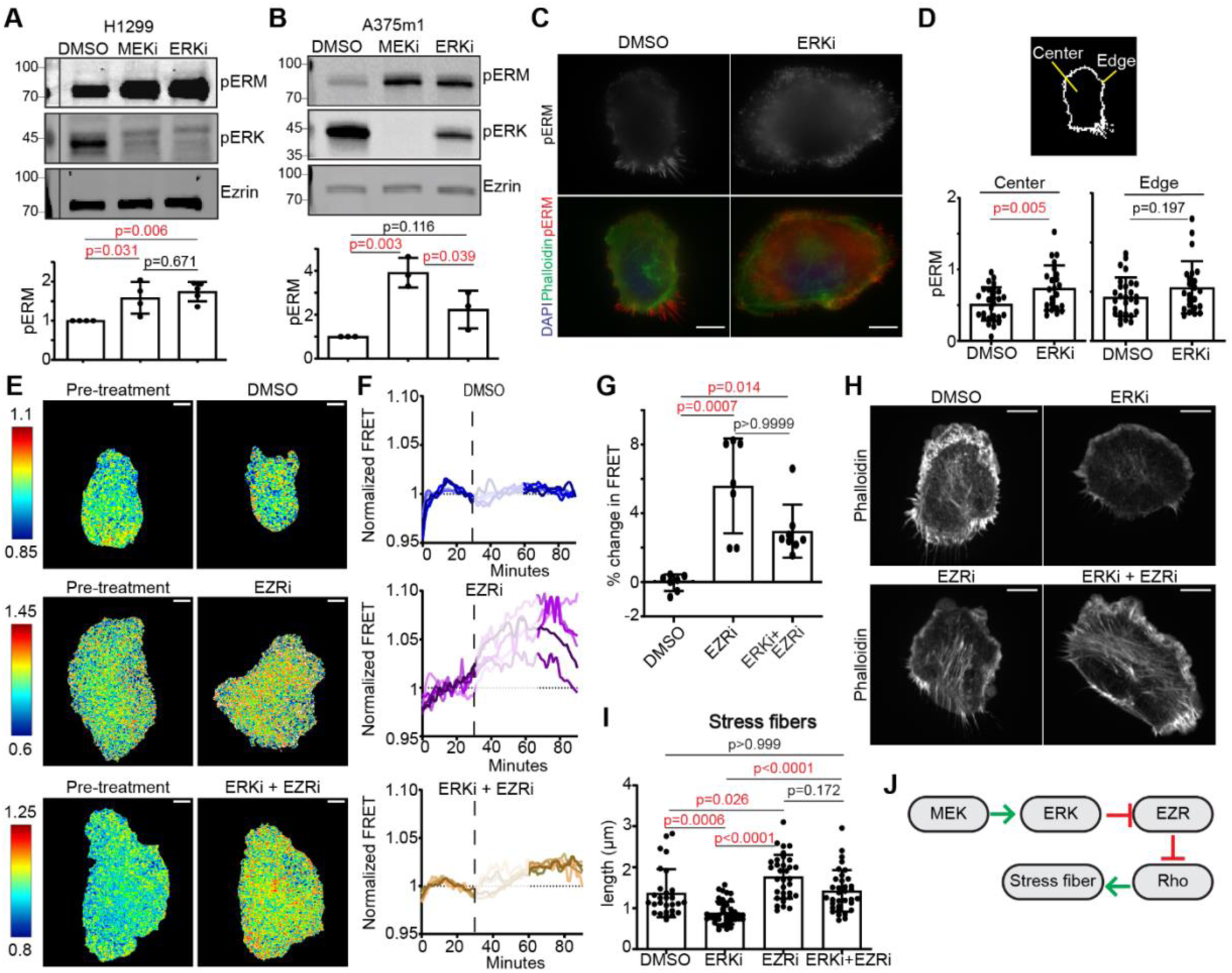
ERK inhibits Ezrin, which suppresses Rho activity and stress fiber formation. (A, B) Representative Western blot and quantification of phospho-ERM (pERM) T567, normalized to total Ezrin in cells treated with DMSO, MEKi (AZD6244 5 µM), or ERKi (SCH772984 5 µM). (C) Representative epifluorescence images of pERM immunofluorescence in H1299 cells treated with DMSO (n=28) or ERKi (n=23). (D) Quantification of pERM in cell center and edge. (E) Representative Rho activity heatmaps (F) and time courses in H1299 cells treated with DMSO (n=7), Ezrin I (EZRi, NSC668394 2.5 µM; n=7), or ERKi+EZRi (n=8); dashed line indicates time of treatment, and FRET ratios are normalized to mean pre-treatment level. (G) Quantification of the change in FRET ratio in E. (H, I) Phalloidin staining of H1299 cells treated with DMSO (n=29), ERKi (n=42), EZRi (n=33), or ERKi+EZRi (n=34) and quantification of stress fiber length in the cell body. (J) Schematic of ERK-mediated regulation of Ezrin, Rho activity, and stress fiber formation. Scale bars, 10 µm. Significance by one-way ANOVA (A–C, G, I) or Student’s t-test (D).

We tested the model that ERK inhibits Ezrin to induce Rho and stress fibers by testing if Ezrin inhibition phenocopies active ERK. Migrating H1299 cells treated with the Ezrin inhibitor NSC668394 exhibited increased Rho activity, measured by Rho-GTP pulldown assays and Rho2G FRET (Figures 2E-G and S2E). Cells co-treated with both ERK and Ezrin inhibitors also exhibited elevated RhoA activity, supporting a model in which Ezrin acts downstream of ERK (Figures 2E-G). Similarly, Ezrin inhibition increased the length of stress fibers in the cell body, while the stress fibers in cells with both ERK and Ezrin inhibition were indistinguishable from those under Ezrin inhibition alone (Figures 2H and 2I). Thus, ERK promotes stress fiber length primarily by reversing Ezrin’s inhibition of RhoA (Figure 2J).

### ERK phosphorylates LOK Thr952 to inhibit Ezrin

Since LOK (lymphocyte-oriented kinase/ serine threonine kinase 10, STK10) is the major ERM kinase(*54*), we tested if ERK regulation of Ezrin involves LOK. We generated CRISPR-Cas9 knockouts of LOK in H1299 cells with two independent guide RNAs targeting exon 1 and control cells with a guide RNA targeting the nonresident protein GFP (Figure S3A). Clonal cell lines established from each target showed comparable reductions in cell migration velocity (Figures S3B-D). Using gRNA-1 LOK knockout (LOK-KO) cells, we found that ERK inhibition did not affect pERM levels in the absence of LOK, indicating that ERK’s regulation of Ezrin requires LOK (Figure 3A). Previous phosphoproteomics studies have identified LOK Thr952 as a direct ERK substrate. The phosphosite (PNPSpTPSKA) conforms to the canonical ERK consensus motif (PXpS/TP), is sensitive to RAF, MEK, and ERK inhibitors, and is directly phosphorylated by an ATP-analog-sensitive ERK2 mutant (*55*). In order to validate that ERK phosphorylates LOK T952 during cell migration, we raised a phospho-specific antibody against LOK pT952 (pLOK). Using 293T cells expressing S-tagged LOK, western blotting with anti-phospho-LOK T952 revealed a clear band at ∼110 kDa, consistent with the molecular weight of LOK, overlapping with the total S-LOK protein, and sensitive to dephosphorylation with calf intestinal phosphatase (CIP) (Figure S3E). The antibody failed to recognize phospho-deficient LOK T952A mutant (Figure S3F). We used the pLOK antibody to confirm the endogenous site is controlled by ERK. In 293T cells, expression of constitutively active BRAF^V600E^ increased LOK T952 phosphorylation, while the inactive MEK AA (S218A/S222A) had no effect and MEK^DD^ (S218D/S222D) trended to increase LOK T952 phosphorylation (Figure S3G). Treating the BRAF^V600E^ and MEK^DD^ cells with MEK or ERK inhibitor reversed LOK T952 phosphorylation to basal levels (Figure 3B). MEK and ERK inhibition similarly decreased LOK T952 phosphorylation H1299 and A375m1 cells (Figures 3C and 3D).

**Figure 3.**
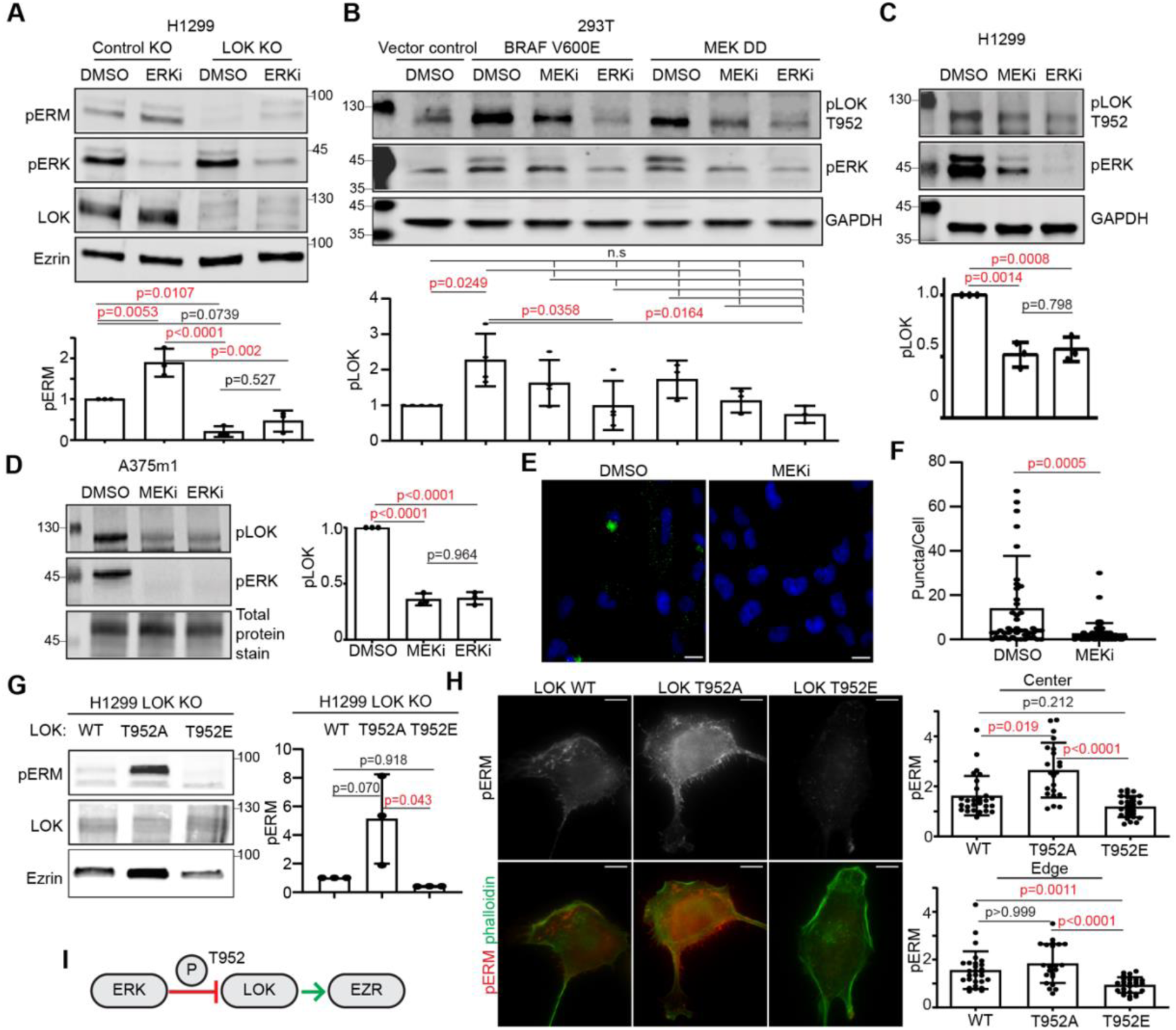
ERK phosphorylates LOK at T952 to inhibit Ezrin activation. (A) Western blot and quantification of pERM T567, normalized to total Ezrin in H1299-GFP KO and H1299-LOK KO (guide RNA1) cells. (B) Western blot and quantification of phospho-LOK (pLOK) T952, normalized to GAPDH, in 293T cells transfected with BRAF^V600E^ and MEK^DD^. (C) Western blot of pLOK T952 in H1299 and (D) A375m1 cells treated with DMSO, MEKi (5 µM, 1 h), or ERKi (5 µM, 1 h). (C, D) Quantification of pLOK normalized to GAPDH or total protein stain. (E, F) Representative image of proximity ligation assay (PLA) for pERK and LOK-WT-S-tag in H1299 cells treated with DMSO (n=46) or MEKi (5 µM; n=60) and DAPI stain of nuclei. Quantification is PLA puncta (green) per cell. Data are from two independent experiments. Scale bars, 100 µm. (G) Western blot and quantification of pERM T567, normalized to Ezrin, in H1299 LOK KO cells stably expressing LOK-WT, LOK-T952A, or LOK-T952E. (H) Representative immunofluorescence and quantification of pERM T567 in H1299 cells stably expressing LOK-WT (n=26), LOK-T952A (n=24), or LOK-T952E (n=27), from two independent experiments. Scale bars, 10 µm. (I) Schematic of ERK-mediated phosphorylation of LOK at T952 inhibiting Ezrin activation. Significance by one-way ANOVA (A–D, G, H) or Student’s t-test (F).

We hypothesized that ERK inhibits Ezrin in the cell body to activate Rho through phosphorylation of LOK T952. To test this, we first tested if ERK interacted with LOK in the cell body, by proximity ligation assay (PLA) with antibodies against phospho-ERK (pERK) and the S-tag of transiently transfected S-tagged LOK. Indeed, control cells treated with DMSO exhibited PLA fluorescent puncta distributed throughout the cell body and excluded from the cell edge and nucleus (Figures 3E and 3F). Treatment with MEK inhibitor reduced the number of PLA puncta (Figures 3E and 3F). We then tested if LOK T952 phosphorylation controls Ezrin activation by stably re-introducing LOK T952 phosphosite mutants into the H1299 LOK KO cells. Cell populations expressing wild-type LOK (LOK WT), a phospho-deficient mutant (LOK T952A), or a phospho-mimetic mutant (LOK T952E) were sorted to obtain similar expression levels of reintroduced LOK. Western blotting and immunofluorescence for pERM showed that the LOK T952A cells harbored increased pEzrin and LOK T952E-expressing cells harbored reduced pEzrin, compared to LOK WT rescue cells (Figure 3G and 3H). Furthermore, the increase in pEzrin in LOK T952A cells, compared to LOK WT cells, specifically occurred in the cell body (Figure 3H). Thus, these data on ERK and LOK support a model in which ERK phosphorylates LOK at T952 in the cell body, where ERK inhibits Ezrin (Figure 3I). Since the LOK T952 phosphosite is on LOK’s flexible C-terminal tail, which is involved in LOK autoinhibition and interaction with Ezrin(*56*) but has not been resolved within the LOK structure, predictions of how the phosphorylation affects LOK’s ability to phosphorylate Ezrin are not readily available.

### ERK controls Rho, stress fibers, and cell morphology through LOK T952 phosphorylation

We tested if ERK controls Rho activity through LOK phosphorylation using the LOK mutant cell lines. Rho-GTP pulldown assays showed that LOK T952A cells harbored reduced Rho-GTP and LOK T952E cells harbored increased Rho-GTP, compared to LOK WT controls (Figure 4A). Imaging the Rho2G biosensor in these cells showed that in the presence of LOK T952E, ERK inhibition no longer regulated cell body RhoA activity, demonstrating that ERK-mediated regulation of RhoA depends on LOK T952 phosphorylation (Figures 4B, 4C, and S4A).

**Figure 4:**
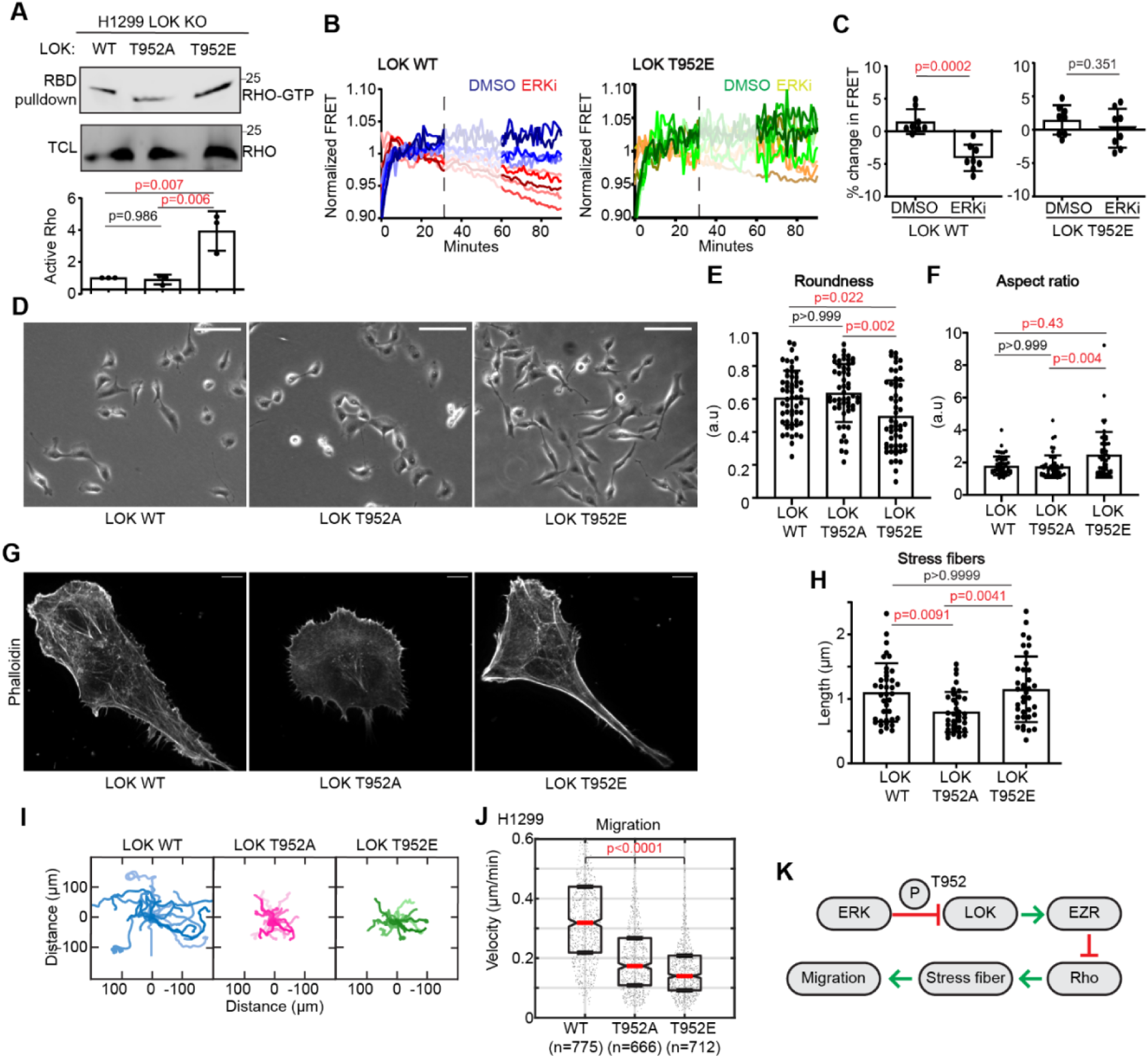
ERK-induced stress fiber formation and cell elongation require LOK T952 phosphorylation. (A) Western blot and quantification of active RHO-GTP from RBD-pulldown, normalized to total RHO in TCL in LOK KO cells with stable LOK-WT, LOK-T952A, and LOK-T952E expression. (B) Representative Rho activity time course in LOK-WT and LOK-T952E cells treated with DMSO or ERKi (WT DMSO n=8; ERKi, n=8. T952E DMSO n=8; ERKi, n=8). (C) Quantification of changes in FRET ratio from B. (D) Representative images of LOK-WT (n=55), LOK-T952A (n=53), and LOK-T952E (n=59) cells. (E, F) Quantification of roundness and aspect ratio from D. Scale bars, 100 μm. (G) Representative phalloidin staining of LOK-WT (n=38), LOK-T952A (n=37), and LOK-T952E (n=37) cells. (H) Quantification of stress fiber length in the cell body from G. Scale bars, 10 μm. (I, J) Rose plots and quantifications of migrating LOK-WT, LOK-T952A, and LOK-T952E cells. (K) Schematic of ERK signaling through LOK–Ezrin to regulate Rho activation, stress fiber formation, and cell migration. Significance by one-way ANOVA (A, E, F, H), Student’s t-test (C), or Kolmogorov–Smirnov test (J).

LOK T952A cells appeared highly spread with large lamellipodia, quantified as increased cell roundedness and a lower aspect ratio (Figures 4D-F). In contrast, LOK T952E cells exhibited elongated morphology with indistinct protrusions, quantified as decreased roundedness and a higher aspect ratio (Figures 4D-F). The elongated phenotype of LOK T952E cells is consistent with hyperactivation of Rho(*57*, *58*). Phalloidin staining of the actin cytoskeletal showed that LOK T952A cells displayed extensive actin transverse arcs associated with lamellipodial protrusion, whereas LOK T952E cells lacked transverse arcs and instead harbored a network of ventral stress fibers and cortical actin (Figures 4G and 4H). Both LOK T952A and LOK T952E mutations reduced migration velocity and persistance, compared to cells expressing LOK WT (Figures 4I, 4J and S4B). This suggests that cell migration requires an optimal balance of LOK T952 phosphorylation and Ezrin inhibition to obtain effective RhoA activation (Figure 4K).

We interrogated the putative ERK/LOK/Ezrin/ARHGAP18/Rho pathway by attempting to rescue the migration defects of the LOK phosphosite mutants. Since inhibition of ERK and LOK T952 phosphorylation results in elevated Ezrin activity, we treated the LOK T952A cells with low dose Ezrin inhibitor (2µM). This level of Ezrin inhibition did not affect LOK WT cell migration, but slightly increased LOK T952A migration velocity and persistance (Figures 5A, 5B, and S5A). Since ERK activation and LOK phosphorylation inhibits Ezrin and ARHGAP18 recruitment to the cell body, we tested if over-expression of ARHGAP18 could rescue the LOK T952E migration defect. Indeed, co-expressing ARHGAP18 with LOK T952E increased migration, compared to cells expressing LOK T952E alone, while the migration of cells co-expressing ARHGAP18 with LOK WT was not different from that of cells expression LOK WT alone (Figure 5C, 5D and S5B). Lastly, since Rho induces stress fiber formation through Rho-associated kinase (ROCK)(*30*), we tested the LOK inhibition of Rho signal by treating LOK T952E cells with a low dose ROCK inhibitor (ROCKi, Y-27632, 3µM). While this level of ROCKi did not affect LOK WT cell migration velocity, it increased LOK T952E migration velocity (Figures 5E, 5F and S5C). Thus, the slowed migration in LOK T952A cells is partly due to overactive Ezrin and the slowed migration of LOK T952E cells is partly due to depleted ARHGAP18 and excessive RhoA-ROCK signaling.

**Figure 5:**
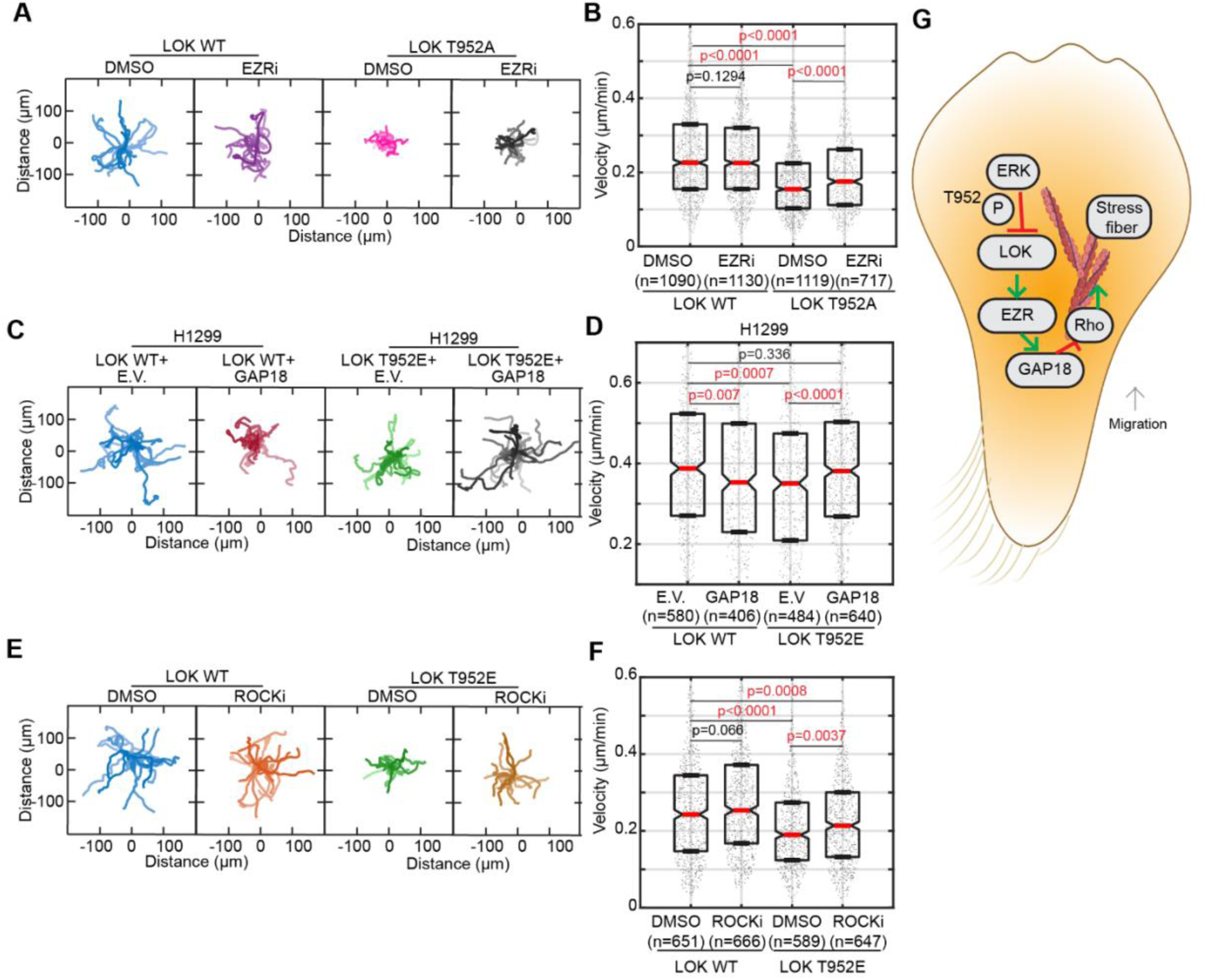
Modulation of Ezrin, ARHGAP18, and ROCK rescue migration defects in LOK T952 mutant cells. (A, B) Rose plots and quantification of migrating LOK-WT and LOK-T952A cells treated with DMSO or EZRi (NSC668394 2 μM). (C, D) Rose plots and quantification of migrating H1299 cells expressing LOK-WT (with empty vector or ARHGAP18) and LOK-T952E (with empty vector or ARHGAP18). (E, F) Rose plots and quantification of migrating LOK-WT and LOK-T952E cells treated with DMSO or ROCKi (Y-27632, 3μM). Significance tested by Kolmogorov–Smirnov test. (G) Schematic of ERK/LOK/Ezrin/ARHGAP18/Rho signaling model, which promotes stress fiber formation in the cell body to drive migration.

## Discussion

This study reveals the mechanism behind ERK’s induction of Rho activity and contractile stress fibers needed for cell migration: ERK phosphorylates LOK at T952, which inhibits LOK’s activation of Ezrin and releases Ezrin’s inhibition of RhoA in the cell body. In this way, LOK T952 phosphorylation promotes the assembly of contractile actin stress fibers and cell migration. No other proposed mechanism for ERK regulation of Rho explains how ERK induces Rho and contractile stress fiber assembly specifically in the cell body. We and others show that ERK phosphorylates LOK T952 in a range of mouse and human cells, including fibroblast and cancer cells known to move with mesenchymal mode migration, suggesting that this signal is a fundamental mechanism of mammalian cell migration control(*55*).

While Ezrin and the ERM protein family canonically function as membrane-actin linkers for the actin cortex (*44*), other reports are consistent with our conclusion that Ezrin inhibits Rho signaling within the cell body. The interaction of ERMs’ N-terminal FERM domain with phosphatidyl inositol 4,5 bisphosphate (PIP2) in the membrane promotes ERM activation by releasing an intramolecular inhibition with the ERM C-terminus (*44*, *56*, *59*). However, new computational modeling suggests that Ezrin’s intramolecular inhibition can also be spontaneously released and T567 phosphorylation impedes re-association (*60*). Ezrin’s FERM domain binds ARHGAP18 when Ezrin is in a partially open conformation as Ezrin T567E (*45*), consistent with the hypothesis that ERMs exist in multiple conformation states with distinct binding partners (*61*).

We propose that cell body-localized LOK maintains Ezrin in a sufficiently open conformation for an Ezrin FERM-ARHGAP18 interaction and ERK inhibits LOK’s ability to induce this “sufficiently open” Ezrin state. This is supported by our PLA and western blotting, which showed that ERK interacts with LOK throughout the cell body and that ERK inhibition and the LOK T952A mutant increase the proportion of T567-phosphorylated Ezrin. Our findings that Ezrin inhibition and ARHGAP18 knockdown induce cell body RhoA activation and stress fiber assembly are consistent with Ezrin and ARHGAP18 functioning together to inhibit cell body RhoA during cell migration, similar to Ezrin’s local inhibition of Rho in mammalian intestinal epithelial cells by recruiting and activating of ARHGAP18 (*45*, *49*). Since ERK does not appear to control the activation of Ezrin or RhoA at the cell edge or in edge protrusions, we conclude that the PIP2-mediated opening of Ezrin at the cell cortex creates a dominant signal for membrane-localized LOK to phosphorylate and activate Ezrin, even in the presence of local ERK activity.

The elucidated ERK/LOK/Ezrin/ARHGAP18/Rho pathway explains how elevated ERK signaling instructs mesenchymal-mode cell migration in response to oncogenic RAS of RAF mutations and growth factor, extracellular matrix, and mechanical cues. Imaging of Rho biosensors showed that ERK regulates baseline Rho activity in the cell body during migration and controls the formation of ventral stress fibers populated with myosin II, which assemble and contract near the basal membrane (*22*). The regulation of baseline Rho activity differs from ERK’s control of Rho activity pulses in migrating cell sheets (*13*), likely due to the sustained increase in ERK activity in H1299 cells with RAS mutation and the lack of a directional cue in random walk migration (*9*). RAS pathway oncogenes and high growth factor stiumulation shift spontaneous ERK activity pulses (∼ every 20-30 minutes) to increased basal ERK activity or merged, sustained pulses (*62–64*). Thus, we propose that at any given time, most cells in an activated population will harbor high levels of active ERK and pLOK, low Ezrin activity, and high RhoA activity for migration. ERK induces protrusion–retraction cycles at the leading edge (*65*, *66*) and new adhesions that form during edge protrusion will amplify ERK activity in protrusions (*67*, *68*), which further drives actin assembly and adhesion turnover (*12*, *65*, *69–72*) and also diffuses to the cell body where it promotes stress fiber assembly and actomyosin contractility to pull the cell body forward. Instantaneous, feedback-induced drops in ERK activity result in un-phosphorylated LOK, which activates spontaneously-opened Ezrin molecules in the cell body, leading to Ezrin interaction with ARGHGAP18 and transient declines in RhoA activity and pauses in motion.

## Methods

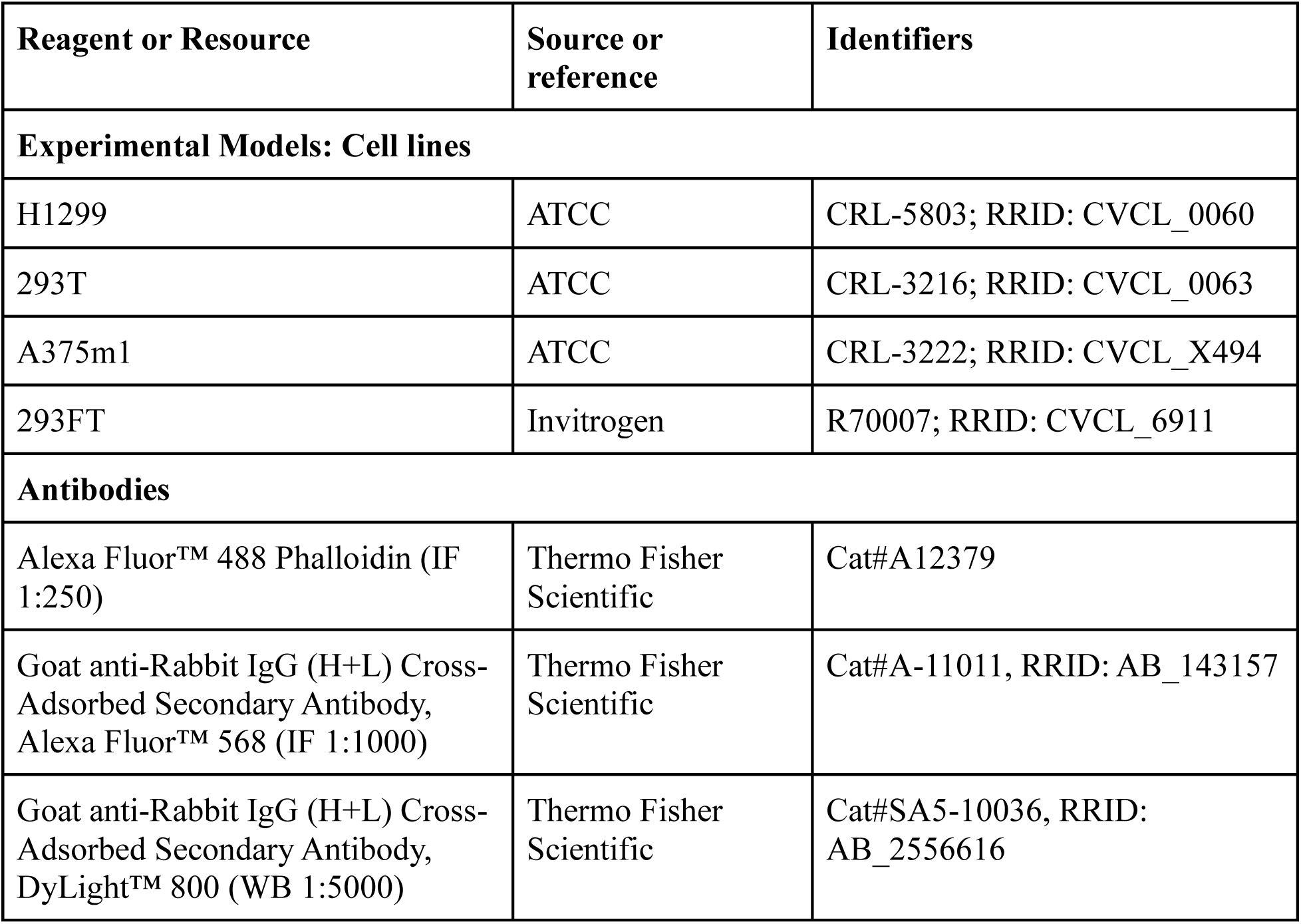
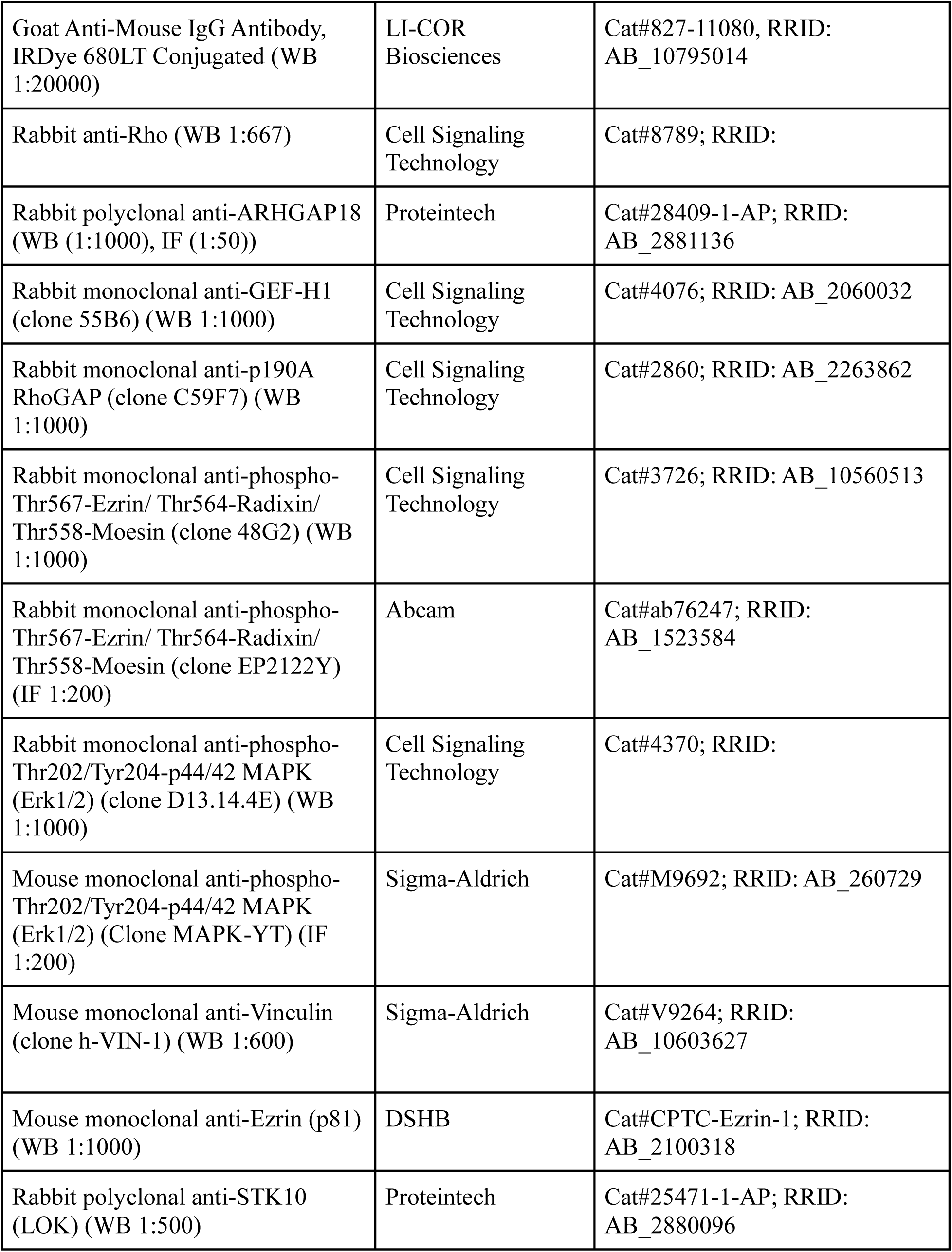
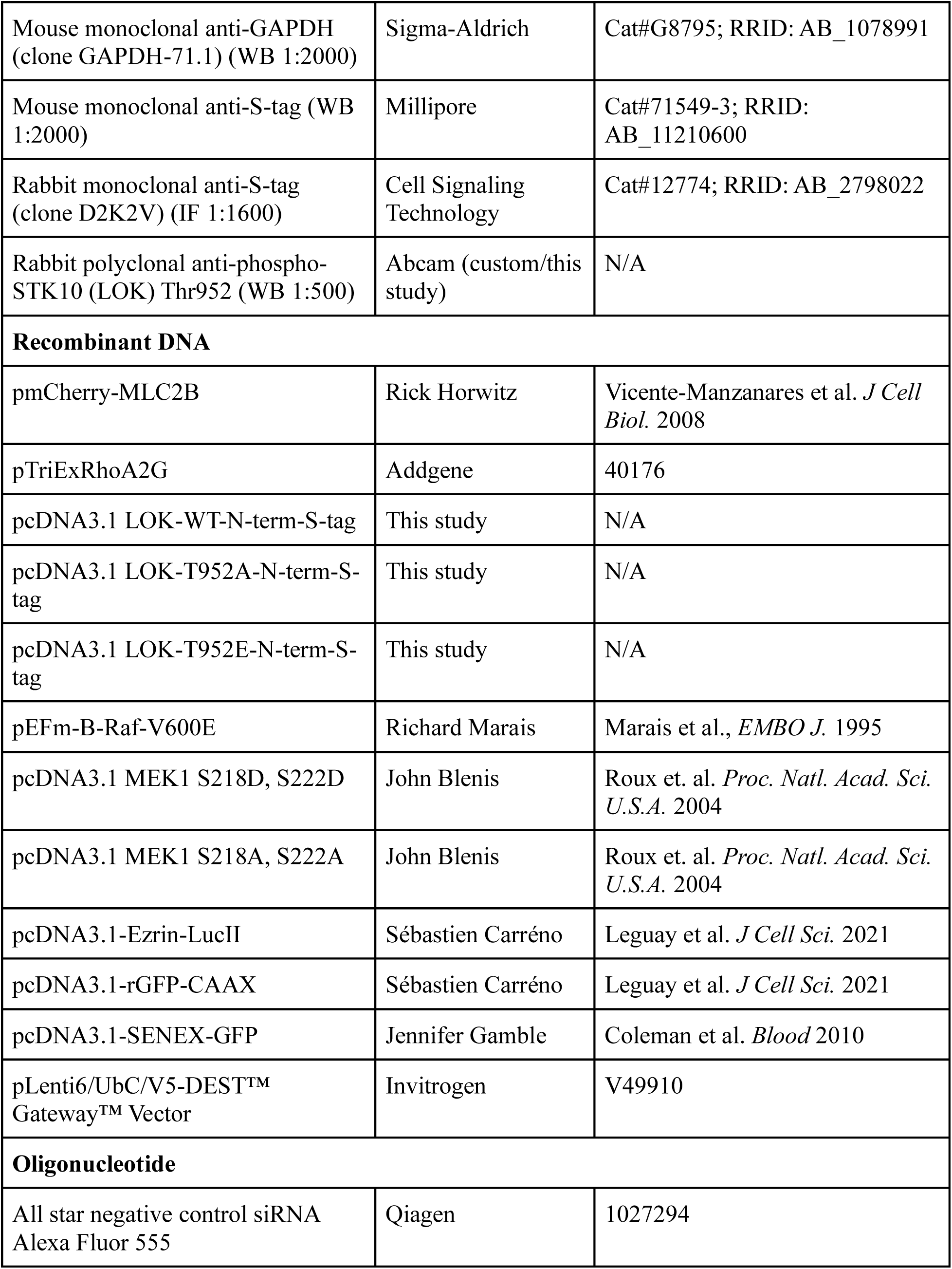
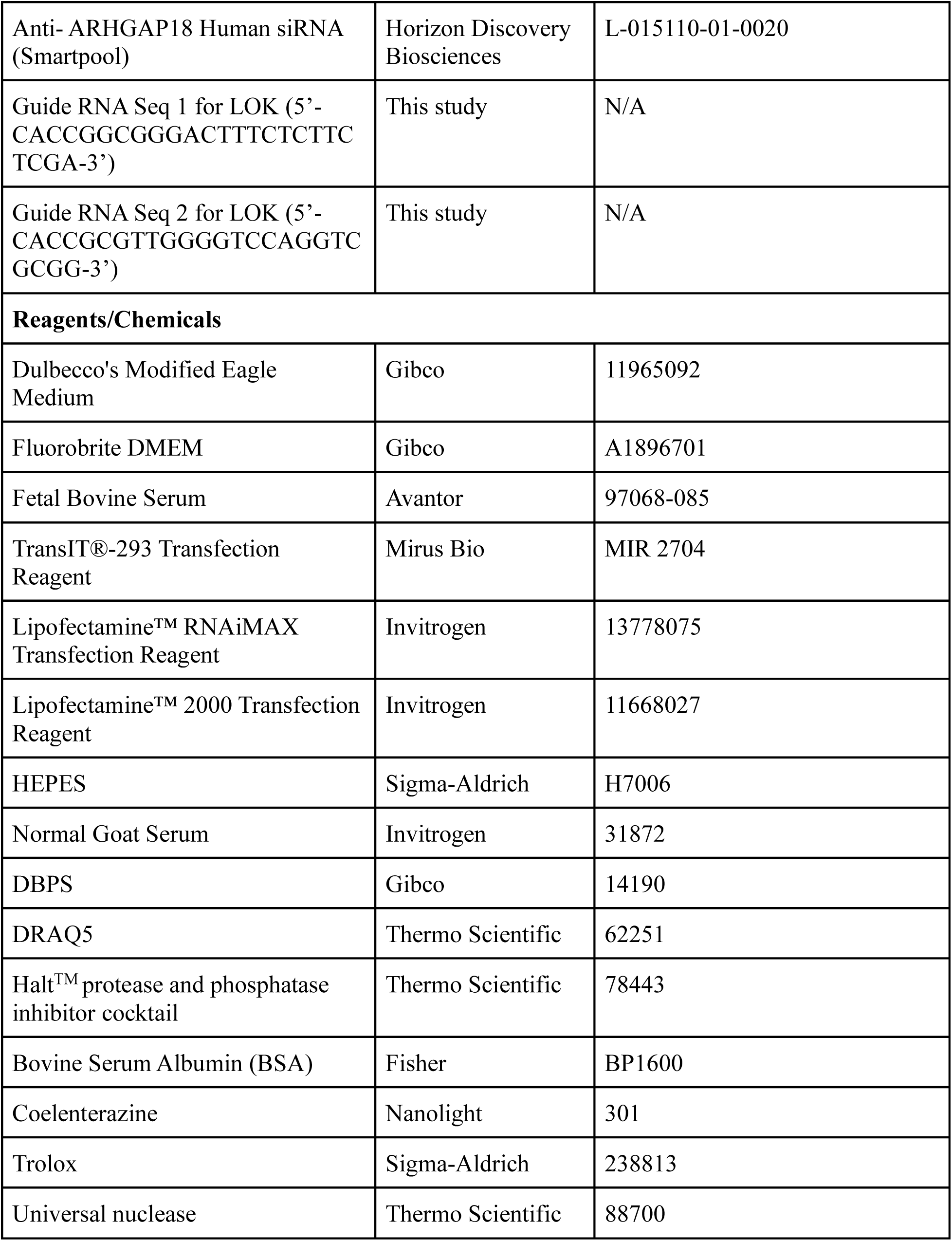
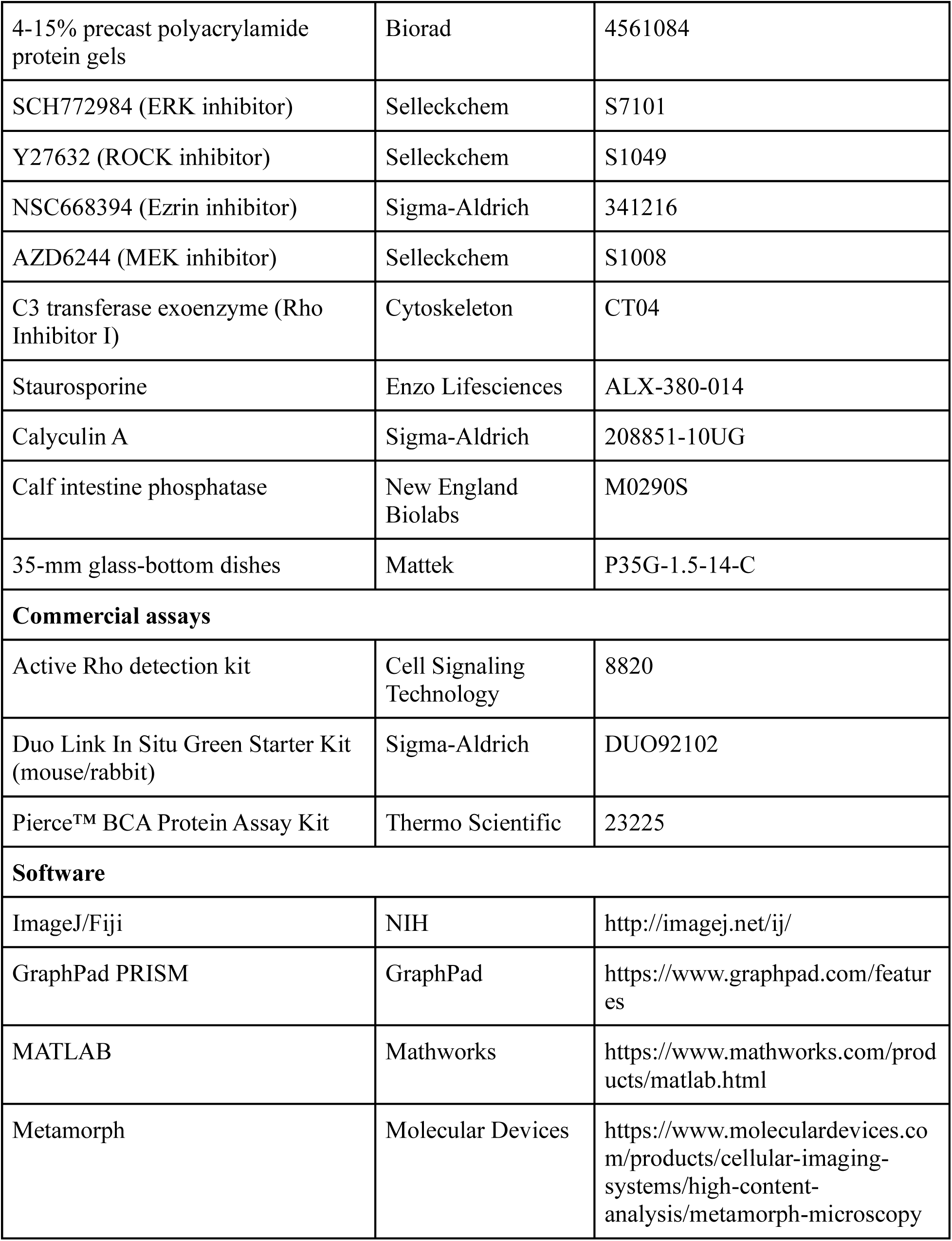
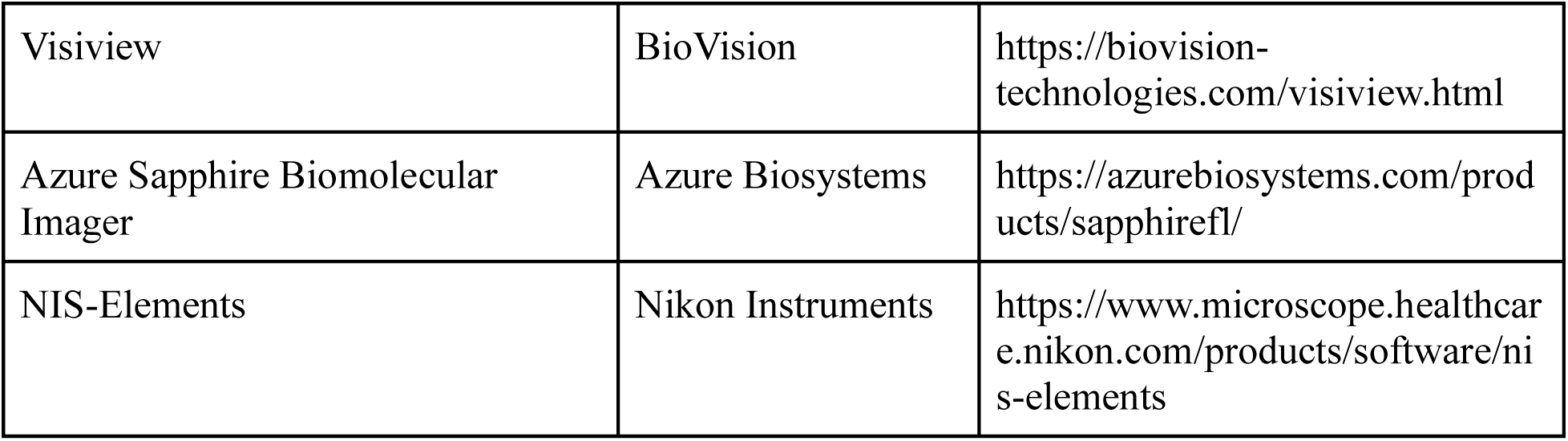

### Tissue culture and cell line generation

Cell lines were maintained at subconfluent density in DMEM with 5% FBS at 37 °C and 5% CO₂. Transient plasmid transfection was performed using TransIT-293 for HEK293T cells and Lipofectamine 2000 for H1299 cells, following the manufacturer’s instructions. siRNA transfections were by RNAiMAX according to the manufacturer’s protocol. siRNAs were co-transfected with fluorescently labeled non-targeting Allstar control siRNA (Sigma) to assess transfection efficiency. >95% of cells were positive for the fluorescent signal. BRAF^V600E^, MEK, ERK, and Ezrin inhibitor treatments were 1 hr before assay. Rho inhibitor treatment was 4 hours before assay.

RhoA2G stable H1299 cell lines were generated by transfection with pTriEx-RhoA2G followed by multiple rounds of fluorescence-activated cell sorting. LOK knockout (KO) H1299 cell lines were generated using Cas9/CRISPR-mediated genome editing. Cells were transfected with Cas9 and guide RNAs targeting GFP the LOK genomic sequence (guide RNA seq 1and 2), selected with puromycin (2 μg/mL), and cloned by limiting dilution. WT H1299 cells were transfected with a GFP-targeting CRISPR/Cas9 construct and selected in parallel as a negative control. LOK deletion was confirmed by Sanger sequencing. For stable, lentivirus LOK re-introduction, LOK WT, T952A, and T952E constructs with N-terminal mScarlet and C-terminal S-tag were cloned into pLenti6/UbC/V5-DEST Gateway vector using Gibson assembly. Lentiviral particles were generated in 293FT cells using TransIT-293 transfection with the pLenti6 lentiviral transfer construct and packaging plasmids (p59, p60, p61). H1299 LOK knockout cells (guide RNA 1) were infected with viral supernatants in the presence of polybrene (4 µg/ml) twice. Stable cell lines expressing LOK-mScarlett constructs were sorted by FACS selection using the Texas-Red channel.

### Immunofluorescence staining

Cells were grown on 12-well glass-bottom dishes, fixed with 4% paraformaldehyde in PHEM buffer (60 mM PIPES, 25 mM HEPES, 10 mM EGTA, 4 mM MgSO₄, 50 mM β-glycerophosphate, pH 6.9) supplemented with 1 mM sodium orthovanadate for 20 min at room temperature, and washed with PBS. For MLC-mCherry staining, prior to fixation, cells were pre-extracted for 45–60 s with buffer containing 0.2% Triton X-100, 4% PEG-8000, 5 mM NaCl, 100 mM PIPES, 1 mM EGTA, and 1 mM MgCl₂. Cells were then permeabilized with 1% CHAPS in PHEM buffer for 5 min at room temperature and washed in PHEM buffer. For F-actin staining, cells were incubated with Alexa Fluor 488–conjugated phalloidin (1:250, Invitrogen) in PBS with 1% BSA for 20 min at room temperature, washed with PBS, and imaged with 1 mM Trolox antifade reagent. For phospho-ERM and ARHGAP18 staining, cells were blocked with 10% normal goat serum (NGS) in MBST (50 mM MOPS, 150 mM NaCl, 0.05% Tween-20, pH 7.4) for 1 h at room temperature before incubation with anti-phospho-Ezrin and anti-ARHGAP18 in MBST with 5% NGS overnight at 4 °C. Cells were then washed with MBST, incubated with Alexa Fluor 561–conjugated secondary antibodies, Alexa Fluor 488 phalloidin, and DAPI in MBST with 5% NGS for 1 h at room temperature, washed with MBST and PBS, and imaged in PBS with 1 mM Trolox.

### Microscopy of fixed cells

Confocal images of ARHGAP18-, MLC-, phospho-ERM-, duolink proximity ligation and phalloidin were acquired on a Nikon Ti inverted microscope with a Yokogawa CSU-10 spinning disk confocal (Borealis modification, Spectral Applied Research), 405/488/561 nm solid-state lasers, and a Photometrics Myo CCD camera, using MetaMorph software. Epifluorescence images of phospho-ERM were collected on a Nikon Ti inverted microscope with a CFI Apo TIRF 60×/1.45 NA oil objective, Perfect Focus, and a Photometrics Prime BSI sCMOS camera.

### Analysis of fixed images

For stress fiber quantification, in MATLAB, cell masks were generated from the 488 nm channel and a cell body sub-mask was created by applying a 5 µm inward offset from the cell boundary. Stress fibers were identified by hysteresis thresholding of the sub-mask and fibermetric (Frangi filter) with parameters: StructureSensitivity = 200, Thickness = 1:10. Fibers were then binarized and objects < 10 pixels excluded. Fiber lengths were calculated from fiber skeletons with distance transforms.

Colocalization between actin fibers and myosin light chain was quantified using the Coloc2 plugin in Fiji (ImageJ) on whole-cell regions of interest (ROIs). The Costes automatic threshold regression method was applied and colocalization assessed by Pearson’s correlation coefficients (PCC). A point spread function (PSF) of 2.0 was used, and significance was determined using Costes randomization with 10 iterations.

For phospho-Ezrin/ARHGAP18 intensity quantification, fluorescence intensities were quantified from the 561 nm channel. Cells were masked and a cell center sub-mask was created by applying a 1 µm inward offset from the cell boundary for phospho-Ezrin quantification and 5 µm inward offset for ARHGAP18 quantification. Total fluorescence intensity within the sub-mask was summed and normalized to the sub-mask area to obtain the mean fluorescence intensity per unit area.

### Rho–GTP pulldown assay

Active Rho was isolated from sub-confluent cell lysates using the Active Rho Detection Kit, according to the manufacturer’s protocol. Cells were treated with inhibitors for 1 h before lysis. Pulldown eluates and total lysates were mixed with tricine sample buffer (Bio-Rad), boiled, resolved on the same SDS–polyacrylamide gel, transferred to nitrocellulose membrane, and probed with anti-RhoA antibody. Band intensities were quantified in ImageJ, using rectangular regions of interest drawn around each band and background subtracted from an adjacent membrane region.

### RhoA FRET imaging and analysis

Cells expressing pTriExRhoA2G were plated subconfluently in 35-mm glass-bottom dishes. 1 h before imaging, medium was changed to FluoroBrite DMEM with 10% FBS and 20 mM HEPES. Migrating cells were imaged by epifluorescence on a Nikon Ti inverted microscope with a CFI Apo TIRF 60×/1.45 NA oil objective, perfect focus, and a Photometrics Prime BSI scientific CMOS camera at gain 1, 2×2 binning, and 100 MHz readout speed. Illumination was by a Sola SE light engine (Lumencor) at 20% output, with 100 ms exposure. Time-lapse CFP/YFP FRET image pairs were acquired every 20 s for 90 min to capture donor and FRET emission signals, respectively. CFP signals were collected using the Chroma 49001 filter set (excitation 436/20 nm, emission 480/40 nm) with a T455LP dichroic mirror, while FRET/YFP signals were collected using the Chroma 49003 filter set (excitation 436/20 nm, emission 535/30 nm) with a T515LP dichroic mirror. At 30 min, DMSO or inhibitors diluted in FluoroBrite were added using an syringe.

FRET efficiency was quantified using the MATLAB Biosensor Processing Package. Donor and FRET images were background corrected, and pixelwise FRET/CFP ratios were calculated to generate activation maps. Mean cellular FRET values were obtained by averaging all pixels within a cell mask. For spatiotemporal analysis, time-lapse movies of single cells were imported into MATLAB, and cell masks were generated at each timepoint. The cell centroid was tracked over time to define XY trajectories and velocity vectors. A line perpendicular to the instantaneous velocity vector and passing through the centroid defined the cell midline, dividing the cell into “front” and “rear.” Pixels in the rear quartile were used to quantify back-specific RhoA activity.

### Western blotting

Sub-confluent cells were washed with ice-cold PBS and lysed in SDS lysis buffer (1% SDS, 1 mM EDTA, 125 mM Tris-HCl pH 7.4, 2 mM PMSF, Halt protease and phosphatase inhibitor cocktail, 1 μM Pepstatin A, 1 mM DTT, and Universal nuclease). Lysates were normalized to total protein concentration, quantified using the Pierce™ BCA Protein Assay Kit, according to the manufacturer’s instructions, and mixed with Laemmli sample buffer (62.5 mM Tris-HCl pH 6.8, 2% SDS, 10% glycerol, 50 mM DTT, and 0.01% bromophenol blue). Then boiled for 5 min., and resolved on 4–15% precast SDS–polyacrylamide gels (Bio-Rad) in Tris–glycine–SDS running buffer (25 mM Tris base, 192 mM glycine, 0.1% SDS, pH 8.3) and transferred to nitrocellulose membranes in Tris–glycine transfer buffer (25 mM Tris base, 192 mM glycine, 20% methanol, pH 8.3). Membranes were blocked in 5% BSA in TBST (20 mM Tris-HCl pH 7.5, 150 mM NaCl, 0.1% Tween-20) for 1 h at room temperature, incubated with primary antibodies in 5% BSA/TBST overnight at 4 °C, and then with Alexa Fluor–conjugated secondary antibodies (685 nm or 784 nm) for 1 h at room temperature.

The membranes were imaged on an Azure Sapphire Biomolecular Imager (Azure Biosystems) at 100 μm resolution using the 685 nm and 784 nm channels (laser power level 3). Band intensities were quantified in ImageJ (NIH). Identical rectangular regions of interest were drawn around each band, background intensity was subtracted from a nearby area of the same membrane, and signals normalized to the corresponding loading control (GAPDH and vinculin). Phosphorylated ERM proteins were normalized to the total Ezrin level. Phospho-LOK was normalized to GAPDH, as both the phospho-LOK and total LOK antibodies were generated in rabbit.

### BRET measurement

H1299 cells were transfected with MyBP-Ezrin-LucII and rGFP-CAAX and seeded into 96-well glass-bottom plates for 24 hr. Cells were then washed with FluoroBrite DMEM media with 10% FBS and 20 mM HEPES, and then incubated with 5 μM coelenterazine for 5 min at room temperature. BRET was measured on a Tecan Infinite 200 PRO multifunctional microplate reader (Tecan), with donor emission at 360–440 nm and acceptor emission at 475–560 nm. BRET ratios were calculated as the ratio of acceptor to donor emission.

### Proximity ligation assay

Proximity ligation assays were performed using the Duolink In Situ Starter Kit according to the manufacturer’s protocol. Cells were fixed on coverslip dishes with 4% PFA in PHEM buffer (60 mM PIPES, 25 mM HEPES, 10 mM EGTA, 4 mM MgSO₄, 50 mM β-glycerophosphate, pH 6.9) supplemented with 1 mM sodium orthovanadate for 20 mins. Cells were washed with PBS, permeabilized with 1% CHAPS in PHEM buffer for 5 mins at room temperature, washed with PHEM buffer, and blocked with 10% normal goat serum in MBST buffer (50 mM MOPS, 150 mM NaCl, 0.05% Tween-20, pH 7.4) for 1 hour at room temperature. Cells were then incubated overnight at 4 °C with primary antibodies against phospho-ERK and S-tag in 5% normal goat serum in MBST and then incubated with Duolink PLUS and MINUS probes for 1 hour at 37 °C and subjected to probe ligation for 30 mins at 37 °C and amplification for 100 mins at 37 °C. Coverslips were mounted in Duolink mounting medium containing DAPI. Fluorescent PLA signals were imaged in the GFP channel using 488 nm excitation on a confocal microscope. GFP and DAPI channels were merged in ImageJ and PLA puncta were quantified by manually counting.

### Cell morphology quantification

Cells were imaged using a Leica DM IL LED phase-contrast microscope (Leica Microsystems) with a 20× air objective and Clara monochrome CCD camera (Andor). Cell boundaries were manually outlined in Fiji/ImageJ and morphological parameters were calculated using the built-in Analyze > Set Measurements/Measure commands (roundness = 4·area/(π·major_axis²)) and aspect ratio from the ‘Fit Ellipse’ outputs (major/minor).

### Random walk migration assay

Cells were plated subconfluently in 12-well glass-bottom plates in DMEM supplemented with 5% FBS. After 48 h, medium was replaced with FluoroBrite DMEM containing 10% FBS and 2 μM DRAQ5. Cells were imaged on a Nikon Ti inverted microscope equipped with a Plan Fluor ELWD 20× air objective, an environmental chamber maintained at 37 °C with 5% CO₂, and an Andor Clara CCD camera (Model DR-328G-CO1-SIL, Andor) using NIS-Elements software. Images were acquired every 10 min for 12 h. Nuclei were detected via the Cy5 channel, and individual cell movements were tracked over time. Cell tracks were analyzed in MATLAB to calculate migration parameters including velocity, directionality, and persistence. Rose plots show tracks of the 20 cells closest to the median migration speed.

### Statistical analysis

Normality was tested using the Shapiro–Wilk test in GraphPad Prism (v10.4.1). For normally-distributed data (p > 0.05), parametric tests were applied (two-tailed unpaired Student’s *t*-test for pairwise comparisons, or one-way ANOVA with Tukey’s post hoc test for multiple comparisons). For datasets that failed the normality test, non-parametric tests were used (Mann–Whitney U test for two groups, Kruskal–Wallis with Dunn’s correction for multiple groups). Data are presented as mean ± s.d. from three independent experiments unless otherwise specified.

## Acknowledgements

We thank the University of Utah Shared Resources, including Flow Cytometry/Cell Sorting, Mutation Generation and Detection, and Cell Imaging Cores. Additionally, we thank Anthony Bretscher from Weill Institute for Cell & Molecular Biology, Ithaca, NY, USA and Jennifer Gamble from University of Sydney, Australia for providing Ezrin and ARHGAP18 expression constructs. The graphical abstract for this manuscript was created in BioRender (BioRender. 2025. BioRender.com. https://www.biorender.com).

## Author contributions

Conceptualization: A.M.K. and M.C.M; formal analysis: A.M.K., J.S., J.P.B., and K.R.C; investigation: A.M.K., J.S., J.P.B.; methodology: A.M.K., J.S., K.R.C., and M.C.M.; writing – original draft: A.M.K. and M.C.M.; writing – review: all authors; data curation: A.M.K and M.C.M.; sample curation: M.C.M.; resources, supervision, and funding acquisition: M.C.M.

## Data and Materials Availability

### Lead contact

Requests for further information and resources should be directed to, and will be fulfilled by, the lead contact, Michelle Mendoza.

### Materials availability

All unique/stable reagents generated in this study are available from the lead contact with a completed materials transfer agreement.

### Data and code availability

This paper does not report any original sequencing or structural data. All original code has been deposited on GitHub: https://github.com/MendozaLabHCI/AutoCell and https://github.com/MendozaLabHCI/ActinFiberQuantification.

## Supplementary Materials

**Figure S1.**
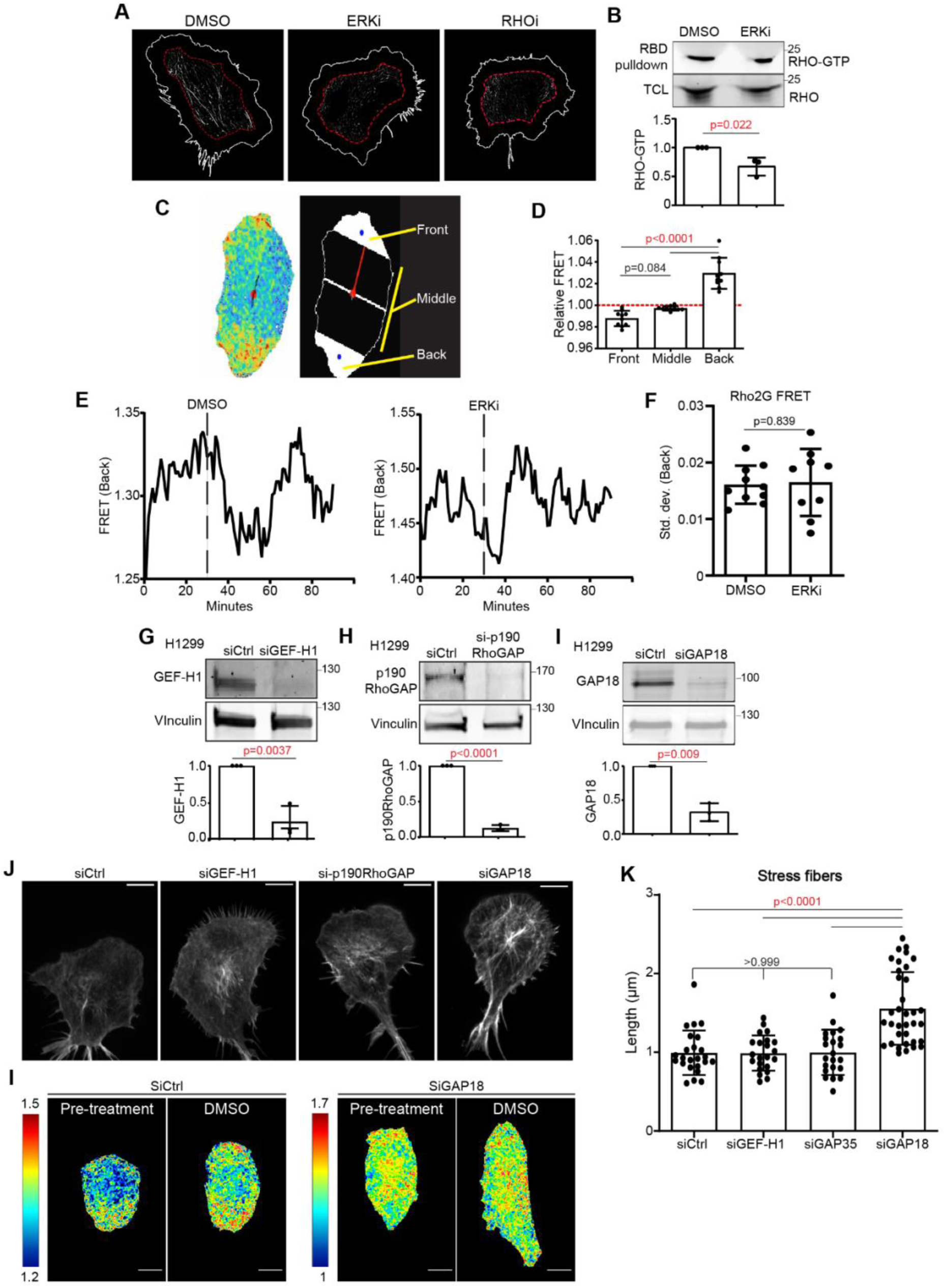
Analysis of stress fibers, Rho activity, and ERK-controlled Rho regulators. (A) Grayscale images of actin fibers in the cell body of representative cells from Fig. 1A; bright fibers correspond to stress fibers. Red dashed lines indicate the cell body region used for stress fiber quantification. (B) Representative western blot and quantification of RHO-GTP from RBD-pulldown, normalized to total RHO in total cell lysates (TCL), from cells treated with DMSO or ERKi (SCH772984, 5 µM). (C) Representative Rho activity heatmap of a single migrating cell (left), schematic of a migrating cell with front, middle, and back regions labeled (right). (D) Control cells, before DMSO treatment, from Fig. 1E. Relative FRET ratio in the front, middle, and back regions, compared to the mean FRET ratio in the total cell (red dashed line). (E) Representative time course of Rho activity at the back of the cell. (F) Quantification of standard deviation of mean Rho activity at the back of the cell after DMSO (n=10) or ERKi (n=9) treatment. (G–I) Representative Western blots and quantification of GEF-H1 (G), p190ARhoGAP (H), and ARHGAP18 (I) following siRNA-mediated knockdown. Quantification is normalized to vinculin. (J, K) Representative phalloidin staining and quantification of stress fiber length in the cell body of cells transfected with siCtrl (n=24), siGEF-H1 (n=22), siP190GAP (n=22), or siGAP18 (n=35). (L) Representative Rho activity heatmaps of migrating cells expressing RhoA2G and transfected with either siCtrl or siARHGAP18. Scale bars, 10 µm. Statistical significance was assessed by Student’s t-test (B, F, G–I) or one-way ANOVA (D, K).

**Figure S2:**
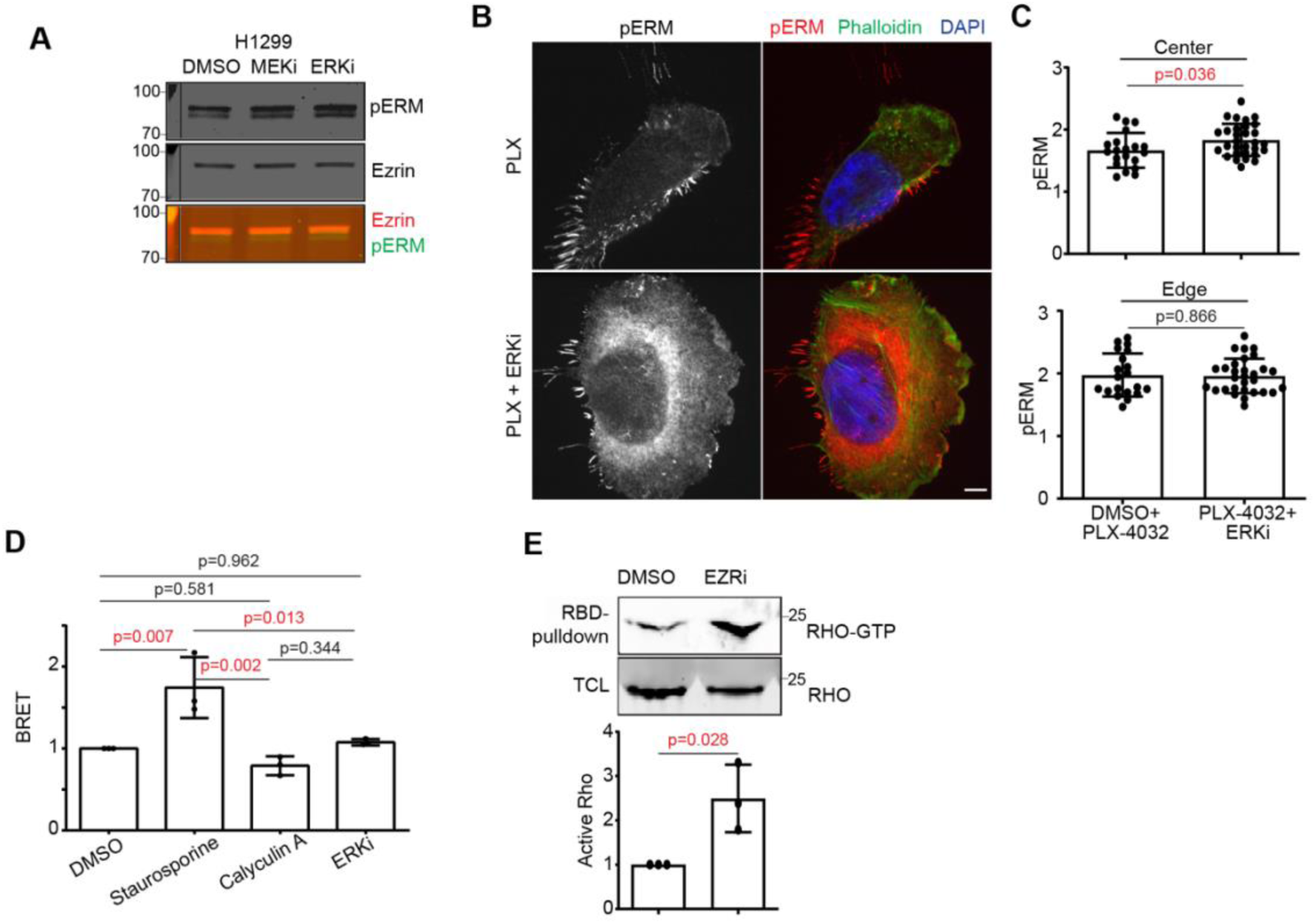
ERK regulates Ezrin phosphorylation and membrane localization. (A) Western blots of pERM phosphorylation T567 in H1299 cells treated with DMSO, MEKi (AZD6244 5 µM), or ERKi (SCH772984 5 µM). The most prominent pERM band (green) overlaps with Ezrin (red). (B) Representative confocal images of immunofluorescence for pERM T567 in H1299 cells treated with 2 µM RAF activator PLX-4032+DMSO (n=21) or 2 µM PLX-4032+5 µM ERKi (n=29). (C) Quantification of pERM intensity in cell center and edge from B. (D) Relative BRET measurement of membrane-bound active ERM in H1299 cells treated with DMSO, staurosporine (100nM, 15min), calyculin A(100nM, 10min), or ERKi (5 µM, 1hr). (E) Representative western blot and quantification of active RHO-GTP from RBD-pulldown, normalized to total RHO in total cell lysates (TCL). EZRi is 2.5 µM NSC668394. Scale bars, 10 µm. Significance by Student’s t-test (C, E) or one-way ANOVA (D).

**Figure S3.**
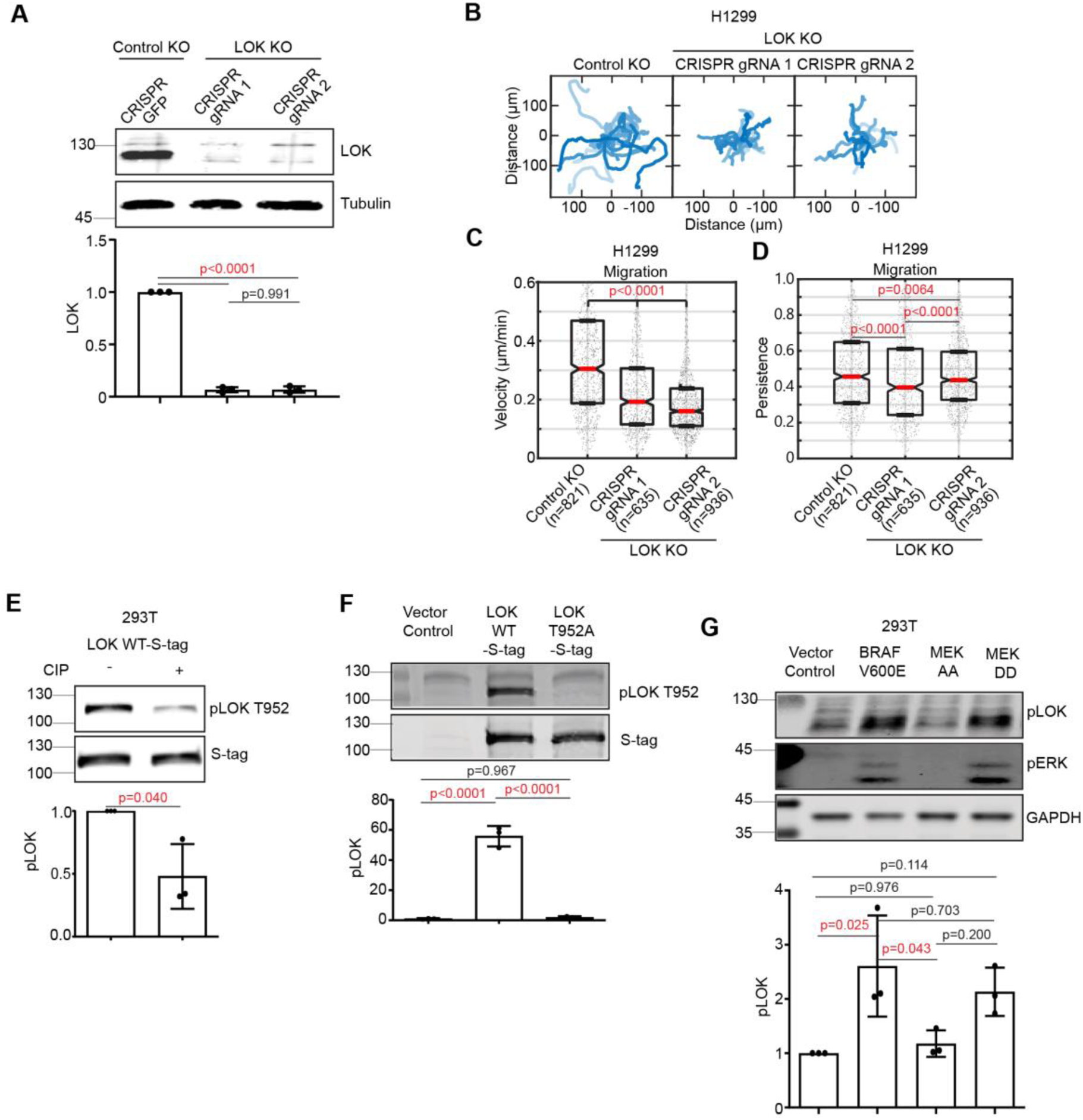
Characterization of LOK knockout and phospho-LOK mutants. (A) Western blot of LOK in H1299 GFP control or LOK knockout clones. LOK quantification is normalized to α-tubulin. (B) Rose plots of migrating H1299 GFP control or LOK knockout clones. Tracks of 20 cells with migration speed closest to the median are shown. (C, D) corresponding quantification of migration velocity and persistence. Significance by Kolmogorov–Smirnov test. (E) Western blot and quantification of LOK T952 phosphorylation (pLOK, normalized to S-tag) in 293T cells transfected with LOK-WT-S-tag and treated with or without calf intestinal phosphatase (CIP). (F) Western blot and quantification of pLOK T952, normalized to total ERK in 293T cells transfected with empty S-tag, LOK-WT-S-tag, or LOK-T952A-S-tag. (G) Western blot and quantification of pLOK T952, normalized to GAPDH in 293T cells transfected with empty vector, BRAF^V600E^, MEK1^AA^, or MEK1^DD^. Significance by one-way ANOVA (A, F, G) and student’s t-test (E).

**Figure S4.**
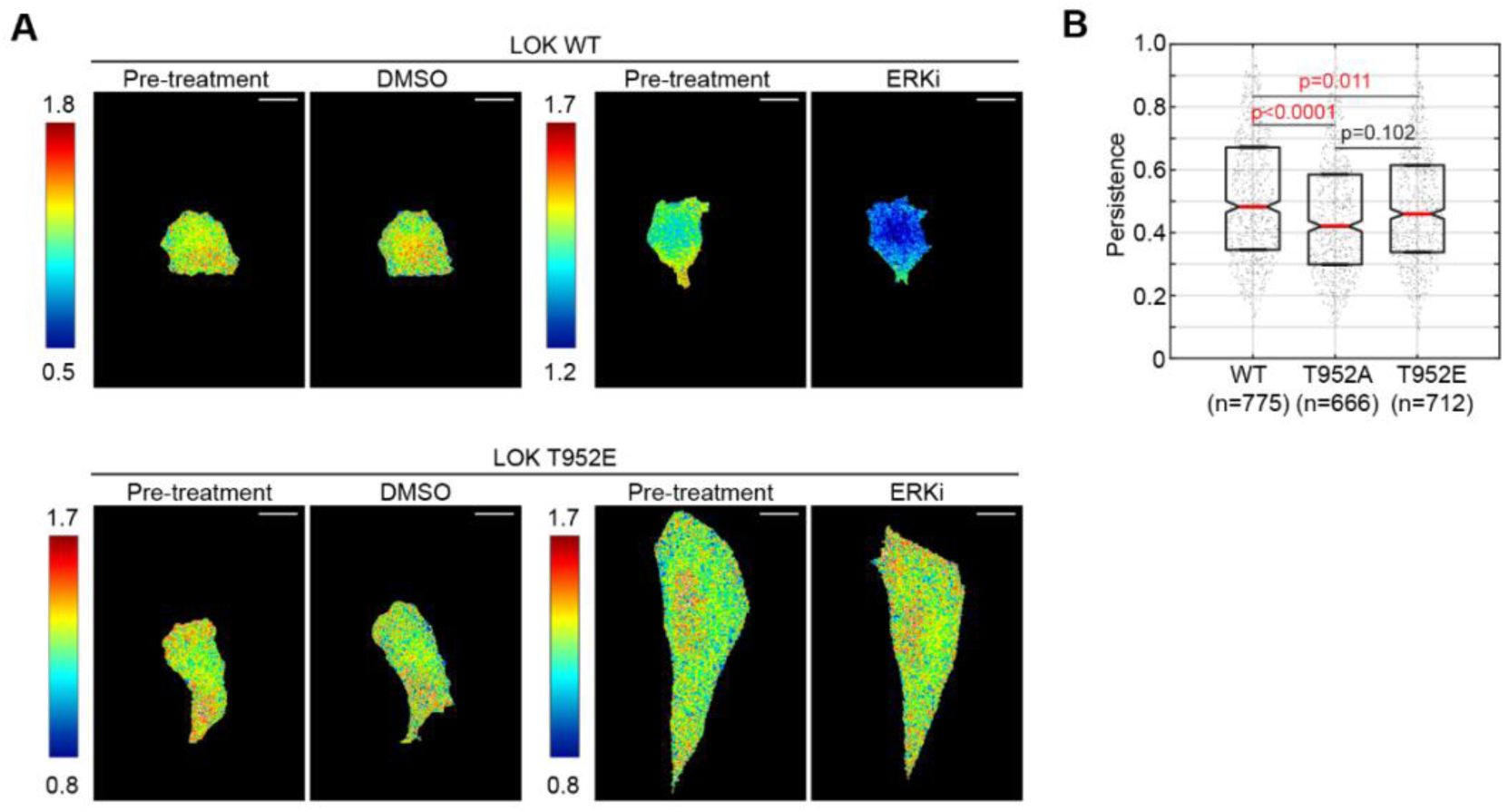
ERK inhibition alters Rho activity and migration persistence in LOK-WT and LOK-T952E cells. (A) Representative Rho activity heatmaps of LOK-WT and LOK-T952E cells treated with DMSO or ERKi. (B) Quantification of migration persistence of cells from Figure 4I and J. Significance by Kolmogorov–Smirnov test.

**Figure S5.**
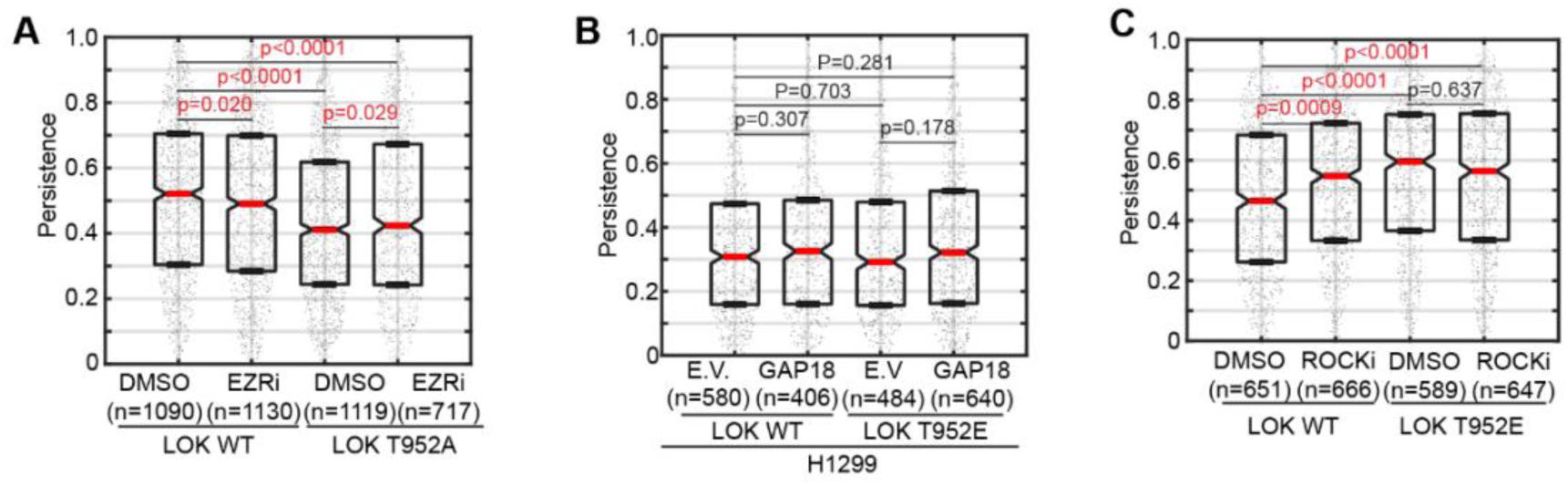
Migration persistence of LOK T952 mutant cells rescued with modulation of Ezrin, ARHGAP18, and ROCK. Quantification of migration persistence of cells from (A) Fig. 5A and B, (B) Fig. 5C and D, and (C) Fig. 5E and F. Significance by Kolmogorov–Smirnov test.

**Supplemental movie 1. Rho activity in a migrating H1299 cell with DMSO added at time point 91. Each timepoint is 20 sec.**

**Supplemental movie 2. Rho activity in a migrating H1299 cell with ERK inhibitor added at at time point 91. Each timepoint is 20 sec.**

## References

1. S. SenGupta, C. A. Parent, J. E. Bear, The principles of directed cell migration. Nat. Rev. Mol. Cell Biol. 22, 529–547 (2021).

2. L. B. Case, C. M. Waterman, Integration of actin dynamics and cell adhesion by a three-dimensional, mechanosensitive molecular clutch. Nat. Cell Biol. 17, 955–963 (2015).

3. P. Agarwal, R. Zaidel-Bar, Principles of Actomyosin Regulation *In Vivo*. Trends Cell Biol. 29, 150–163 (2019).

4. V. Filić, L. Mijanović, D. Putar, A. Talajić, H. Ćetković, I. Weber, Regulation of the Actin Cytoskeleton via Rho GTPase Signalling in Dictyostelium and Mammalian Cells: A Parallel Slalom. Cells 10, 1592 (2021).

5. E. E. Sander, J. P. ten Klooster, S. van Delft, R. A. van der Kammen, J. G. Collard, Rac downregulates Rho activity: reciprocal balance between both GTPases determines cellular morphology and migratory behavior. J. Cell Biol. 147, 1009–1022 (1999).

6. S. Tojkander, G. Gateva, P. Lappalainen, Actin stress fibers – assembly, dynamics and biological roles. J. Cell Sci. 125, 1855–1864 (2012).

7. P. Friedl, Prespecification and plasticity: shifting mechanisms of cell migration. Curr. Opin. Cell Biol. 16, 14–23 (2004).

8. C. D. Lawson, A. J. Ridley, Rho GTPase signaling complexes in cell migration and invasion. J. Cell Biol. 217, 447–457 (2018).

9. S. C. Samson, A. M. Khan, M. C. Mendoza, ERK signaling for cell migration and invasion. Front. Mol. Biosci. 9 (2022).

10. H. Lavoie, J. Gagnon, M. Therrien, ERK signalling: a master regulator of cell behaviour, life and fate. Nat. Rev. Mol. Cell Biol. 21, 607–632 (2020).

11. D. A. Cheresh, J. Leng, R. L. Klemke, Regulation of Cell Contraction and Membrane Ruffling by Distinct Signals in Migratory Cells. J. Cell Biol. 146, 1107–1116 (1999).

12. H. Yu, G. Xiao, M. Gu, L. Zhang, M. Xia, S. Mo, Y. Zhao, C. Wei, pERK transition–induced directional mode switching promotes epithelial tumor cell migration. Proc. Natl. Acad. Sci. 121, e2318871121 (2024).

13. N. Hino, L. Rossetti, A. Marín-Llauradó, K. Aoki, X. Trepat, M. Matsuda, T. Hirashima, ERK-Mediated Mechanochemical Waves Direct Collective Cell Polarization. Dev. Cell 53, 646–660.e8 (2020).

14. K. Aoki, Y. Kondo, H. Naoki, T. Hiratsuka, R. E. Itoh, M. Matsuda, Propagating Wave of ERK Activation Orients Collective Cell Migration. Dev. Cell 43, 305–317.e5 (2017).

15. S. Shin, C. A. Dimitri, S.-O. Yoon, W. Dowdle, J. Blenis, ERK2, but not ERK1, induces epithelial to mesenchymal transformation via DEF motif dependent signaling events. Mol. Cell 38, 114–127 (2010).

16. S. M. Nalluri, C. S. Sankhe, J. W. O’Connor, P. L. Blanchard, J. N. Khouri, S. H. Phan, G. Virgi, E. W. Gomez, Crosstalk between ERK and MRTF-A signaling regulates TGFβ1-induced epithelial-mesenchymal transition. J. Cell. Physiol. 237, 2503–2515 (2022).

17. T. Vallenius, Actin stress fibre subtypes in mesenchymal-migrating cells. Open Biol. 3, 130001 (2013).

18. D. T. Burnette, S. Manley, P. Sengupta, R. Sougrat, M. W. Davidson, B. Kachar, J. Lippincott-Schwartz, A role for actin arcs in the leading-edge advance of migrating cells. Nat. Cell Biol. 13, 371–381 (2011).

19. B. Kovac, J. L. Teo, T. P. Mäkelä, T. Vallenius, Assembly of non-contractile dorsal stress fibers requires α-actinin-1 and Rac1 in migrating and spreading cells. J. Cell Sci. 126, 263–273 (2013).

20. P. Hotulainen, P. Lappalainen, Stress fibers are generated by two distinct actin assembly mechanisms in motile cells. J. Cell Biol. 173, 383–394 (2006).

21. T. Vignaud, C. Copos, C. Leterrier, M. Toro-Nahuelpan, Q. Tseng, J. Mahamid, L. Blanchoin, A. Mogilner, M. Théry, L. Kurzawa, Stress fibres are embedded in a contractile cortical network. Nat. Mater. 20, 410–420 (2021).

22. J. I. Lehtimäki, E. K. Rajakylä, S. Tojkander, P. Lappalainen, Generation of stress fibers through myosin-driven reorganization of the actin cortex. eLife 10, e60710 (2021).

23. D. T. Burnette, L. Shao, C. Ott, A. M. Pasapera, R. S. Fischer, M. A. Baird, C. Der Loughian, H. Delanoe-Ayari, M. J. Paszek, M. W. Davidson, E. Betzig, J. Lippincott-Schwartz, A contractile and counterbalancing adhesion system controls the 3D shape of crawling cells. J. Cell Biol. 205, 83–96 (2014).

24. A. Bolado-Carrancio, O. S. Rukhlenko, E. Nikonova, M. A. Tsyganov, A. Wheeler, A. Garcia-Munoz, W. Kolch, A. von Kriegsheim, B. N. Kholodenko, Periodic propagating waves coordinate RhoGTPase network dynamics at the leading and trailing edges during cell migration. eLife 9, e58165 (2020).

25. F. M. Vega, G. Fruhwirth, T. Ng, A. J. Ridley, RhoA and RhoC have distinct roles in migration and invasion by acting through different targets. J. Cell Biol. 193, 655–665 (2011).

26. J. J. Bravo-Cordero, V. P. Sharma, M. Roh-Johnson, X. Chen, R. Eddy, J. Condeelis, L. Hodgson, Spatial regulation of RhoC activity defines protrusion formation in migrating cells. J. Cell Sci. 126, 3356–3369 (2013).

27. K. Zaoui, S. Honoré, D. Isnardon, D. Braguer, A. Badache, Memo-RhoA-mDia1 signaling controls microtubules, the actin network, and adhesion site formation in migrating cells. J. Cell Biol. 183, 401–408 (2008).

28. M. Nemethova, S. Auinger, J. V. Small, Building the actin cytoskeleton: filopodia contribute to the construction of contractile bundles in the lamella. J. Cell Biol. 180, 1233–1244 (2008).

29. G. Totsukawa, Y. Wu, Y. Sasaki, D. J. Hartshorne, Y. Yamakita, S. Yamashiro, F. Matsumura, Distinct roles of MLCK and ROCK in the regulation of membrane protrusions and focal adhesion dynamics during cell migration of fibroblasts. J. Cell Biol. 164, 427–439 (2004).

30. M. Chrzanowska-Wodnicka, K. Burridge, Rho-stimulated contractility drives the formation of stress fibers and focal adhesions. J. Cell Biol. 133, 1403–1415 (1996).

31. S. Tojkander, G. Gateva, A. Husain, R. Krishnan, P. Lappalainen, Generation of contractile actomyosin bundles depends on mechanosensitive actin filament assembly and disassembly. eLife 4, e06126 (2015).

32. M. L. Kutys, K. M. Yamada, Rho GEFs and GAPs: emerging integrators of extracellular matrix signaling. Small GTPases 6, 16–19 (2015).

33. J. Tong, L. Li, B. Ballermann, Z. Wang, Phosphorylation and Activation of RhoA by ERK in Response to Epidermal Growth Factor Stimulation. PLoS ONE 11, e0147103 (2016).

34. A. von Thun, C. Preisinger, O. Rath, J. P. Schwarz, C. Ward, N. Monsefi, J. Rodríguez, A. Garcia-Munoz, M. Birtwistle, W. Bienvenut, K. I. Anderson, W. Kolch, A. von Kriegsheim, Extracellular Signal-Regulated Kinase Regulates RhoA Activation and Tumor Cell Plasticity by Inhibiting Guanine Exchange Factor H1 Activity. Mol. Cell. Biol. 33, 4526–4537 (2013).

35. S.-H. Fujishiro, S. Tanimura, S. Mure, Y. Kashimoto, K. Watanabe, M. Kohno, ERK1/2 phosphorylate GEF-H1 to enhance its guanine nucleotide exchange activity toward RhoA. Biochem. Biophys. Res. Commun. 368, 162–167 (2008).

36. M. L. Azoitei, J. Noh, D. J. Marston, P. Roudot, C. B. Marshall, T. A. Daugird, S. L. Lisanza, M.-J. Sandí, M. Ikura, J. Sondek, R. Rottapel, K. M. Hahn, G. Danuser, Spatiotemporal dynamics of GEF-H1 activation controlled by microtubule- and Src-mediated pathways. J. Cell Biol. 218, 3077–3097 (2019).

37. J. J. Bravo-Cordero, L. Hodgson, J. S. Condeelis, Spatial regulation of tumor cell protrusions by RhoC. Cell Adhes. Migr. 8, 263–267 (2014).

38. A. Bidaud-Meynard, F. Binamé, V. Lagrée, V. Moreau, Regulation of Rho GTPase activity at the leading edge of migrating cells by p190RhoGAP. Small GTPases 10, 99–110 (2017).

39. A. K. Pullikuth, A. D. Catling, Extracellular Signal-Regulated Kinase Promotes Rho-Dependent Focal Adhesion Formation by Suppressing p190A RhoGAP. Mol. Cell. Biol. 30, 3233–3248 (2010).

40. W. T. Arthur, K. Burridge, RhoA Inactivation by p190RhoGAP Regulates Cell Spreading and Migration by Promoting Membrane Protrusion and Polarity. Mol. Biol. Cell 12, 2711–2720 (2001).

41. W. D. Bradley, S. E. Hernández, J. Settleman, A. J. Koleske, Integrin Signaling through Arg Activates p190RhoGAP by Promoting Its Binding to p120RasGAP and Recruitment to the Membrane. Mol. Biol. Cell 17, 4827–4836 (2006).

42. M. Maeda, H. Hasegawa, T. Hyodo, S. Ito, E. Asano, H. Yuang, K. Funasaka, K. Shimokata, Y. Hasegawa, M. Hamaguchi, T. Senga, ARHGAP18, a GTPase-activating protein for RhoA, controls cell shape, spreading, and motility. Mol. Biol. Cell 22, 3840–3852 (2011).

43. W. R. Thompson, S. S. Yen, G. Uzer, Z. Xie, B. Sen, M. Styner, K. Burridge, J. Rubin, LARG GEF and ARHGAP18 Orchestrate RhoA Activity to Control Mesenchymal Stem Cell Lineage. Bone 107, 172–180 (2018).

44. R. G. Fehon, A. I. McClatchey, A. Bretscher, Organizing the cell cortex: the role of ERM proteins. Nat. Rev. Mol. Cell Biol. 11, 276–287 (2010).

45. A. T. Lombardo, C. A. Mitchell, R. Zaman, D. J. McDermitt, A. Bretscher, ARHGAP18-ezrin functions as an autoregulatory module for RhoA in the assembly of distinct actin-based structures. eLife 13, e83526 (2024).

46. M. Tokunou, T. Niki, Y. Saitoh, H. Imamura, M. Sakamoto, S. Hirohashi, Altered expression of the ERM proteins in lung adenocarcinoma. Lab. Investig. J. Tech. Methods Pathol. 80, 1643–1650 (2000).

47. A.-M. Toms, M. L. Davies, R. Hargest, S. E. Hiscox, W. G. Jiang, Distribution and expression of the ERM family members, ezrin, radixin, moesin and EHM2 in human colon cancer and the clinical relevance. Transl. Gastrointest. Cancer 1, 20514–20214 (2012).

48. K. Kawaguchi, S. Yoshida, R. Hatano, S. Asano, Pathophysiological Roles of Ezrin/Radixin/Moesin Proteins. Biol. Pharm. Bull. 40, 381–390 (2017).

49. R. Zaman, A. Lombardo, C. Sauvanet, R. Viswanatha, V. Awad, L. E.-R. Bonomo, D. McDermitt, A. Bretscher, Effector-mediated ERM activation locally inhibits RhoA activity to shape the apical cell domain. J. Cell Biol. 220, e202007146 (2021).

50. T. Thoresen, M. Lenz, M. L. Gardel, Thick Filament Length and Isoform Composition Determine Self-Organized Contractile Units in Actomyosin Bundles. Biophys. J. 104, 655–665 (2013).

51. R. D. Fritz, M. Letzelter, A. Reimann, K. Martin, L. Fusco, L. Ritsma, B. Ponsioen, E. Fluri, S. Schulte-Merker, J. van Rheenen, O. Pertz, A versatile toolkit to produce sensitive FRET biosensors to visualize signaling in time and space. Sci. Signal. 6, rs12 (2013).

52. W. Qian, N. Yamaguchi, P. Lis, M. Cammer, H. Knaut, Pulses of RhoA signaling stimulate actin polymerization and flow in protrusions to drive collective cell migration. Curr. Biol. CB 34, 245–259.e8 (2024).

53. K. Leguay, B. Decelle, Y. Y. He, A. Pagniez, M. Hogue, H. Kobayashi, C. Le Gouill, M. Bouvier, S. Carréno, Development of conformational BRET biosensors that monitor ezrin, radixin and moesin activation in real time. J. Cell Sci. 134, jcs255307 (2021).

54. N. V. Belkina, Y. Liu, J.-J. Hao, H. Karasuyama, S. Shaw, LOK is a major ERM kinase in resting lymphocytes and regulates cytoskeletal rearrangement through ERM phosphorylation. Proc. Natl. Acad. Sci. 106, 4707–4712 (2009).

55. E. B. Ünal, F. Uhlitz, N. Blüthgen, A compendium of ERK targets. FEBS Lett. 591, 2607–2615 (2017).

56. T. Pelaseyed, R. Viswanatha, C. Sauvanet, J. J. Filter, M. L. Goldberg, A. Bretscher, Ezrin activation by LOK phosphorylation involves a PIP2-dependent wedge mechanism. eLife 6, e22759 (2017).

57. H. F. Paterson, A. J. Self, M. D. Garrett, I. Just, K. Aktories, A. Hall, Microinjection of recombinant p21rho induces rapid changes in cell morphology. J. Cell Biol. 111, 1001–1007 (1990).

58. A. J. Ridley, A. Hall, The small GTP-binding protein rho regulates the assembly of focal adhesions and actin stress fibers in response to growth factors. Cell 70, 389–399 (1992).

59. S. Bosk, J. A. Braunger, V. Gerke, C. Steinem, Activation of F-Actin Binding Capacity of Ezrin: Synergism of PIP2 Interaction and Phosphorylation. Biophys. J. 100, 1708–1717 (2011).

60. D. Vasiliauskas, J. Beiter, S. S. Iyer, A. T. Lombardo, M. C. Mendoza, G. A. Voth, On the Mechanism of Ezrin Activation. bioRxiv [Preprint] (2025). 10.1101/2025.11.07.687285.

61. R. Viswanatha, J. Wayt, P. Y. Ohouo, M. B. Smolka, A. Bretscher, Interactome Analysis Reveals Ezrin Can Adopt Multiple Conformational States. J. Biol. Chem. 288, 35437–35451 (2013).

62. L. J. Bugaj, A. J. Sabnis, A. Mitchell, J. E. Garbarino, J. E. Toettcher, T. G. Bivona, W. A. Lim, Cancer mutations and targeted drugs can disrupt dynamic signal encoding by the Ras-Erk pathway. Science 361, eaao3048 (2018).

63. T. J. Aikin, A. F. Peterson, M. J. Pokrass, H. R. Clark, S. Regot, MAPK activity dynamics regulate non-cell autonomous effects of oncogene expression. eLife 9, e60541 (2020).

64. T. E. Gillies, M. Pargett, J. M. Silva, C. K. Teragawa, F. McCormick, J. G. Albeck, Oncogenic mutant RAS signaling activity is rescaled by the ERK/MAPK pathway. Mol. Syst. Biol. 16, e9518 (2020).

65. M. C. Mendoza, M. Vilela, J. E. Juarez, J. Blenis, G. Danuser, ERK reinforces actin polymerization to power persistent edge protrusion during motility. Sci. Signal. 8, ra47 (2015).

66. M. C. Mendoza, Phosphoregulation of the WAVE regulatory complex and signal integration. Semin. Cell Dev. Biol. 24, 272–279 (2013).

67. H. Yu, G. Xiao, M. Gu, L. Zhang, M. Xia, S. Mo, Y. Zhao, C. Wei, pERK transition–induced directional mode switching promotes epithelial tumor cell migration. Proc. Natl. Acad. Sci. 121, e2318871121 (2024).

68. N. Hino, K. Matsuda, Y. Jikko, G. Maryu, K. Sakai, R. Imamura, S. Tsukiji, K. Aoki, K. Terai, T. Hirashima, X. Trepat, M. Matsuda, A feedback loop between lamellipodial extension and HGF-ERK signaling specifies leader cells during collective cell migration. Dev. Cell 57, 2290–2304.e7 (2022).

69. M. C. Mendoza, E. E. Er, W. Zhang, B. A. Ballif, H. L. Elliott, G. Danuser, J. Blenis, ERK-MAPK drives lamellipodia protrusion by activating the WAVE2 regulatory complex. Mol. Cell 41, 661–671 (2011).

70. D. J. Webb, K. Donais, L. A. Whitmore, S. M. Thomas, C. E. Turner, J. T. Parsons, A. F. Horwitz, FAK-Src signalling through paxillin, ERK and MLCK regulates adhesion disassembly. Nat. Cell Biol. 6, 154–161 (2004).

71. S. Radtke, M. Milanovic, C. Rossé, M. De Rycker, S. Lachmann, A. Hibbert, S. Kermorgant, P. J. Parker, ERK2 but not ERK1 mediates HGF-induced motility in non-small cell lung carcinoma cell lines. J. Cell Sci. 126, 2381–2391 (2013).

72. J.-M. Yang, S. Bhattacharya, H. West-Foyle, C.-F. Hung, T.-C. Wu, P. A. Iglesias, C.-H. Huang, Integrating chemical and mechanical signals through dynamic coupling between cellular protrusions and pulsed ERK activation. Nat. Commun. 9, 4673 (2018).

